# The multifaceted role of acetamide derivative of Chalcone: Anti-inflammatory Action and Impact on Osteoclastogenesis, insights on NF-κB and MAPK pathways

**DOI:** 10.64898/2026.03.20.713114

**Authors:** Sobia Anjum, Tazeem Akram, Unnatti Sharma, Omesh Manhas, Jasha Momo H. Anal, Gurleen Kour, Zabeer Ahmed

## Abstract

Inflammation serves as a vital physiological process essential for preserving health and countering illness. Yet, persistent inflammation drives osteoclastogenesis and ongoing bone erosion in rheumatoid arthritis (RA), mainly via macrophage activation and overproduction of pro-inflammatory cytokines like TNF-α, IL-1β, and IL-6. Limitations of prolonged conventional treatments underscore the need for safer small-molecule inhibitors that address both inflammation and osteoclast formation. Chalcones, natural plant defense compounds, exhibit diverse pharmacological properties including anti-inflammatory, anticancer, antibacterial, antifungal, and antiparasitic actions, owing to their characteristic reactive α, β- unsaturated carbonyl moiety. This study assessed chalcone derivative 7a for its anti-inflammatory effects *in vitro* and *in vivo*, alongside its capacity to modulate osteoclast differentiation, offering the inaugural demonstration of its dual anti-inflammatory and anti-osteoclastogenic properties. In LPS-stimulated macrophages, 7a substantially curtailed nitric oxide production, curbed pro-inflammatory cytokines (TNF-α, IL-1β, IL-6), and concentration-dependently diminished iNOS and COX-2 expression while inhibiting reactive oxygen species levels. *In vivo*, oral 7a dosing potently alleviated carrageenan-evoked paw swelling and restored serum lactate dehydrogenase and C-reactive protein to normalcy. In LPS-exposed mice, it further lowered systemic cytokines and rectified dysregulated biomarkers such as LDH, ALP, ALT, AST, creatinine, and urea. Moreover, in RANKL-stimulated osteoclast cultures, 7a markedly suppressed osteoclastogenesis by downregulating pivotal markers like tartrate-resistant acid phosphatase (TRAP) and matrix metalloproteinase-9 (MMP-9). Derivative 7a also enhances antioxidant defense—superoxide dismutase and catalase—via blockade of NF-κB and MAPK pathways. Overall, chalcone derivative 7a displays robust anti-inflammatory and anti-osteoclastogenic activity, positioning it as a compelling candidate for RA therapy.

## 1. Introduction

Inflammation represents a fundamental physiological response essential for host defense and tissue homeostasis, profoundly influencing disease pathogenesis. Canonical triggers, including infection and tissue injury, activate this process by orchestrating leukocyte recruitment and plasma protein extravasation to the affected site. However, inflammatory pathways can also be activated by a variety of non-infectious stimuli. In particular, para-inflammation—a localized, low-grade inflammatory response, can be triggered by tissue stress or functional disturbance. This adaptive mechanism serves as an intermediary between classical inflammation and basal homeostasis and is primarily mediated by tissue-resident macrophages [1][2]. The innate immune system’s antigen-presenting cells, including macrophages and dendritic cells, initiate the inflammatory cascade upon recognizing pathogen-associated molecular patterns (PAMPs) or damage-associated molecular patterns (DAMPs) [3]. This recognition activates multiple intracellular signaling pathways, most notably the nuclear factor-kappa B (NF-κB) pathway, which regulates the expression of pro-inflammatory genes and facilitates the recruitment of additional immune cells to the site of infection or tissue damage [4]. Moreover, the production of eicosanoids, such as prostaglandins and leukotrienes, via cyclooxygenase-2 and lipoxygenase pathways, plays a key role in the inflammatory process. These eicosanoids modulate vascular permeability, promote vasodilation, and enhance the migration of inflammatory cells, further amplifying the immune response [5]. Furthermore, the reactive oxygen species (ROS) generated during the inflammatory response can exacerbate tissue damage, fueling a self-perpetuating cycle of inflammation that can contribute to chronic disease progression [6]. Pathogenic stimuli, such as lipopolysaccharides (LPS), bind to pattern recognition receptors on immune cells, triggering the release of potent inflammatory mediators, including cytokines and chemokines. This process not only amplifies the local immune response but also enhances the production of acute-phase proteins from the liver, such as C-reactive protein (CRP), which serves as a key biomarker of inflammation [7]. Effective modulation of the immune response is crucial in mitigating the risk of chronic inflammatory diseases, including cardiovascular diseases (CVDs), cancer, and rheumatoid arthritis, highlighting the importance of targeted therapeutic strategies in controlling inflammation and its associated health implications Osteoclasts are multinucleated cells responsible for bone resorption and originate from the monocyte/macrophage lineage [8]. These cells express specific markers, such as tartrate-resistant acid phosphatase (TRAP), and their activity is tightly regulated by various signaling pathways, particularly the receptor activator of nuclear factor-kappa B (RANK) and its ligand, RANKL [9]. The key regulators of osteoclast differentiation are RANKL and macrophage colony-stimulating factor (M-CSF). RANKL, often referred to as the osteoclast differentiation factor, is essential for the initiation and promotion of osteoclastogenesis [10]. Upon the binding of RANKL to its receptor RANK, a cascade of downstream signaling pathways, including NF-kB and MAPK, is triggered. These pathways culminate in the activation of nuclear factor of activated T cells cytoplasmic 1 (NFATc1), an osteoclast-specific transcription factor and the key regulator of osteoclast differentiation [11, 12]. Given its central role in osteoclastogenesis, targeting this signaling axis holds significant promise as a therapeutic strategy for osteolytic inflammatory diseases.

Chalcones are a diverse group of flavonoid ketones characterised by a three-carbon α, β-unsaturated carbonyl group attached to two aromatic rings. These compounds play a pivotal role in plant defence mechanisms, particularly in counter-acting ROS [13]. Chalcone derivatives, such as isoliquiritigenin, flavokawain, and xanthohumol, have demonstrated significant cytotoxicity and the ability to induce apoptosis, thereby promoting anti-tumor activities [14]. The presence of the reactive α, β-unsaturated system within the chalcone structure is responsible for a broad spectrum of pharmacological properties, including enzyme inhibition, anticancer, anti-inflammatory, antibacterial, antifungal, antimalarial, antiprotozoal, and anti-filarial effects. Structural modifications, particularly the introduction of various substituent groups to the aromatic rings, can enhance the compound’s potency, minimise toxicity, and expand its pharmacological potential [15]. The compact structure and Michael acceptor properties of chalcones make them promising candidates for the design of novel derivatives and the exploration of structure-activity relationships. The structure, NMR, and HPLC profile of these chalcone derivatives have already been studied [16].

In this study, we evaluated the anti-inflammatory effects of chalcone derivative 7a in LPS-stimulated RAW264.7 macrophages. Compound 7a suppressed nitric oxide (NO) production, pro-inflammatory cytokines (TNF-α, IL-6, IL-1β), reactive oxygen species (ROS) accumulation, and mRNA expression of iNOS and Cox-2. It also enhanced intracellular levels of superoxide dismutase (SOD) and catalase. Mechanistically, 7a inhibited phosphorylation of key NF-κB (p65, IκBα) and MAPK (p38, ERK) pathway components. These findings were collaborated *in vivo* using carrageenan-induced acute paw edema and LPS-induced systemic inflammation models. Given its promising anti-inflammatory profile, we conducted preliminary *in vitro* anti-arthritic assessments via RANKL-induced osteoclastogenesis model in RAW264.7 cells. 7a markedly reduced the area of tartrate-resistant acid phosphatase (TRAP)-positive multinucleated osteoclasts, RANKL-induced pro-inflammatory cytokines, and MMP-9 levels. It also attenuated ROS production while elevating SOD and catalase levels, as confirmed by ELISA and qRT-PCR. RANKL-induced phosphorylation of p65, IκBα, ERK, and p38 was similarly inhibited. Thus, 7a exhibits a dual therapeutic effect by targeting inflammation and osteoclastogenesis, warranting further investigation of its anti-arthritic potential, including validation in collagen-induced arthritis (CIA) model.

## 2. Materials and Methodology

### 2.1 Materials

The RAW 264.7 cell line used in this study was obtained from the American Type Culture Collection (ATCC). Reagents and solvents were sourced from Sigma-Aldrich, including the anti-inflammatory standard drug Dexamethasone, L-NAME, Griess reagent, 2’, 7’-dichlorodihydrofluorescein diacetate (DCFDA) dye, Lipopolysaccharide (LPS), and Carrageenan lambda. Fetal Bovine Serum (FBS), Dulbecco’s Modified Eagle’s Medium (DMEM), Penicillin-Streptomycin (Penstrap), and ELISA kits for TNF-α, IL-6, IL-1β and MMP-9 were procured from Invitrogen-Gibco (Massachusetts, USA). The MTT dye and Bovine Serum Albumin (BSA) were sourced from HiMedia (India). For RANKL-induced osteoclastogenesis, recombinant RANKL protein was purchased from Thermo Fisher (USA) while the TRAP staining kit was obtained from Takara (China). SOD and Catalase ELISA kits were acquired from G Biosciences (St, Louis, US). Other reagents such as Glycine, Tris-base, Bradford reagent, loading dye, APS, SDS, and bis-acrylamide were purchased from Bio-Rad (California, USA). Antibodies used for Western blotting, including p-NF-κB p65, NF-κB p65, p-IκBα, IκBα, iNOS, Phospho-p44/42 MAPK (ERK1/2), Total-p44/42 MAPK (ERK1/2), p-p38 MAPK, p38 MAPK, β-actin, SOD, Catalase, and anti-rabbit and anti-mouse IgG HRP-conjugated secondary antibodies, were obtained from Cell Signalling Technology (CST, Danvers, MA, USA). PVDF membranes and ECL detection reagents (Immobilon) were also sourced from Cell Signalling Technologies, USA. This collection of reagents and materials enabled the execution of cell culture, inflammatory marker analysis, osteoclastogenesis assays, and *in vivo* inflammation models with precision and reliability.

### 2.2 Studies on LPS induced RAW 264.7 cells

#### 2.2.1 Cell culture

RAW264.7 murine macrophage (ATCC) were maintained in high glucose DMEM, supplemented with 10% FBS, 2.2g/L sodium bicarbonate, sodium pyruvate and 1% penicillin/streptomycin. The cells were sub cultured and maintained at 37°C under 5% CO2 in a humidified incubator.

#### 2.2.2 Cell viability assay

RAW 264.7 cells were seeded in a 96 well plate at a density of 2 × 10^4^ cells per well in 100 µL of DMEM with 10% FBS and allowed to adhere for 24 hours at 37^0^C in a humidified incubator with 5% CO_2_ to assess cell viability of the compounds. The cells were then treated with varying concentrations of eleven chalcone acetamide derivatives (7a-7k) dissolved in DMSO (<0.1%) and incubated for 48 h. Control wells were treated with an equivalent volume of DMEM. After that, 20µL of the MTT solution (2.5mg/mL in 1x PBS) was added to each well, and the plates were incubated for 4 hours to allow the formation of formazan crystals. Finally, 120μl DMSO was added to each well to dissolve formazan crystals formed by the reduction of MTT dye by the mitochondrial enzyme of viable cells. The absorption was recorded at 570 nm wavelength using a microplate reader.

#### 2.2.3 Nitrite estimation

5×10^4^ RAW 264.7 cells were cultured in each well of a 96 well plate. The cells were allowed to acclimatize overnight and then treated with different concentrations of selected chalcone derivative 7a and dexamethasone (positive control) for 1 h. The cells were then stimulated with LPS (1μg/ml) up to 24 h for the release of nitric oxide (NO). NO levels were then detected using Griess reagent and the absorbance was measured at 540 nm. Sodium nitrite was used as standard to confirm the assay’s reliability within the tested concentration range. The half-maximal inhibitory concentration (IC[[) values were calculated by plotting the percentage of NO inhibition versus the concentration of the test compound and fitting the data to a sigmoidal dose-response curve using nonlinear regression analysis.

#### 2.2.4 Cytokine estimation using ELISA

RAW 264.7 cells were cultured in a 96 well plate (5×10^4^ cells/well). The cells were allowed to acclimatize overnight and further treated with different concentrations of the chalcone derivative 7a and dexamethasone (positive control) for 1 h prior to the LPS (1μg/ml) stimulation and incubated for another 24 h. LPS stimulation leads to the production of inflammatory cytokines through the activation of various cellular pathways. After 24h the supernatants were collected and subsequently evaluated for the detection of pro-inflammatory cytokines like TNF-α, IL-6 and IL-1β and anti-inflammatory cytokine IL-10 by using ELISA kits. The absorbance of each sample was compared with the standard curve to measure the cytokine levels.

#### 2.2.5 Measurement of reactive oxygen species

The assessment of LPS triggered intracellular accumulation of ROS was done using fluorescent probe DCFH-DA. Cells were seeded at a density of 5×10^4^ and incubated for 24 hours. After that cells were treated with the different concentrations (1.25μM, 2.5μM and 5μM) of derivative 7a one hour before the stimulation of cells with LPS. After 24h the medium was removed and the cells were washed with PBS. For staining, cells were incubated with 10μM DCFH-DA dye at 37 degrees for 30 min. DCFH- DA is converted to DCFH by the presence of cellular esterase and is further oxidized by the presence of ROS to DCF which is fluorescent and was examined through fluorescent microscopy at 20x magnification.

#### 2.2.6 RT-PCR

RAW 264.7 cells were seeded at a density of 1 × 10[ cells per 60mm petri dish and incubated for 24h. The cells were then treated with the compound 7a at concentrations of 2.5 and 5 µM followed by LPS stimulation for 24 hours at 37°C in a 5% CO[ humidified incubator. Dexamethasone (10 µM) and L-NAME (10 µM) were used as positive controls. Following treatment, total RNA was extracted using Trizol reagent, which was subsequently used for cDNA synthesis with the Promega GoScript^TM^ Reverse Transcription System. Then the quantitative PCR was performed using PowerUp™ SYBR™ Green Master Mix to assess the mRNA expression of inducible nitric oxide synthase (iNOS) and cyclooxygenase-2 (COX-2). qPCR reactions were carried out using the AriaMx Real Time PCR system. The results were expressed as the fold change in target gene expression of treated samples compared to untreated controls, with normalization to the β-actin. The primers used in this study are outlined in Table 1.

#### 2.2.7 Western blotting

RAW 264.7 cells (1 × 10^6^ cells) were seeded in 60mm petri dish and incubated overnight. The cells were then stimulated with LPS and co-incubated with the test compound 7a at concentrations of 2.5 and 5 μM for 24 h at 37 °C in a humidified incubator. Dexamethasone (10μM) was used as a positive control. Total protein was extracted from the cells using an ice-cold cell lysis buffer containing a protease inhibitor cocktail. The protein concentration for each cell lysate was measured using Bradford Protein Assay. For western blotting, an equal amount of protein (30μg) was loaded into each lane of the polyacrylamide gel (4 to 12%) along with a protein ladder. The gel ran into 1x SDS Running Buffer for two hours at 75 Volts. The gel electrophoresis was conducted using the BIO-RAD Mini-PROTEIN Tetra system. After electrophoresis, the protein was electro-transferred to a PVDF membrane through a process which was performed for 120 min at 100 Volts to transfer the protein from the gel to the PVDF membrane. After blocking with 5% BSA for 2 h at room temperature, the membrane was incubated overnight at 4 °C with primary antibodies. The membrane was then washed with 1x TBST and incubated for 1 h at room temperature with HRP conjugated secondary antibody. Proteins were seen with the ECL using ChemiDoc Imaging System (Syngene; Model: G: BOX, chemi XX6). The density of western blotting bands was quantified using the ImageJ software. The density of the protein bands was normalized to that of beta actin. The density values were relatively expressed to the average value for the untreated control group.

### 2.3 *In vivo* study

All experiments were conducted adhering to the guidelines set by the Committee for the Proposal of Control and Supervision of Animal Experiments in each *in vivo* investigation. The institutional animal ethics committee (IAEC) of CSIR-IIIM, Jammu, India provided the approval number (376/84/02/2024) for the studies. BALB/c mice were taken, aged 6–8 weeks and weighing 25-30 g and were housed in polypropylene cages under conventional laboratory settings, which included a 12-hour light/dark cycle, a temperature of 24 ± 2 ◦C, and a relative humidity of 50 ±20%. A regular pellet meal and water were given to the mice. The animals were allowed a week to acclimatise before the experiment.

#### 2.3.1 Carrageenan (CGN) induced paw edema model in BALB/c mice

Carrageenan-induced paw edema is a well-established model for evaluating the anti-inflammatory activity of various compounds. In this model, the administration of carrageenan, a sulfated polysaccharide, leads to the development of acute inflammation in the paw of the animal, characterized by swelling and increased vascular permeability. BALB/c mice were divided into six groups: negative control, positive control, and treatment groups receiving different doses of derivative 7a. The animals were treated with test compound and vehicle, followed by intraplantar injection of carrageenan (50μL of 1% Carrageenan λ prepared in in normal saline). The paw thickness was measured at 0, 2, 4 and 6 h after carrageenan administration using Verniercaliper. The blood samples were also collected for the measurement of CRP and tissue damage marker LDH in the serum. This has been illustrated in Figure 6A.

**Figure 1:**
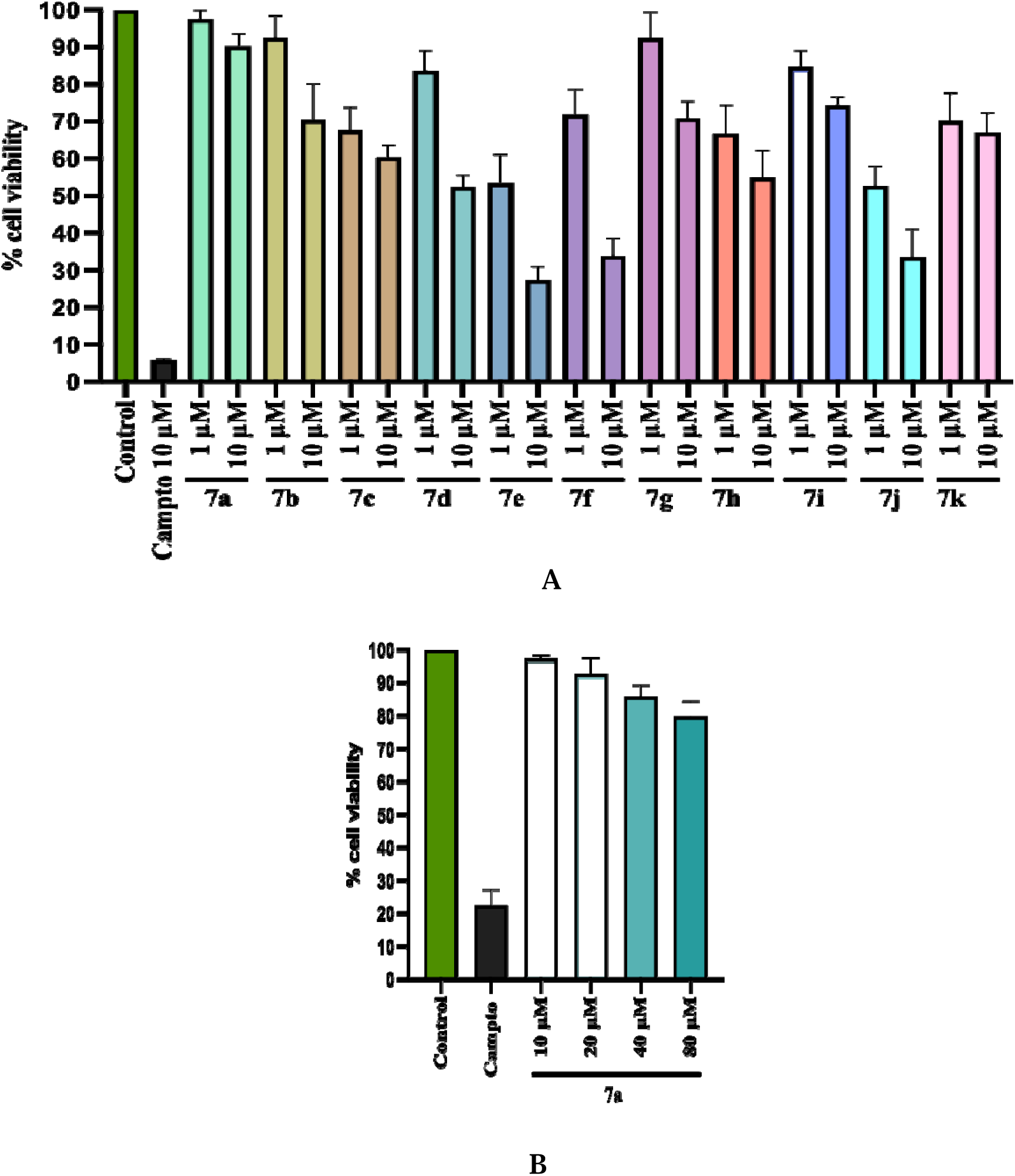
Effect of the chalcone derivatives 7a-7k on percentage cell viability in RAW 264.7 cells at 48h (A) Percentage cell viability of 7a-7k (B) Percentage cell viability of the derivative 7a up to 80μM. The cell viability was determined by MTT assay, the results shown are the mean ± standard deviation of three different experiments.

**Figure 2:**
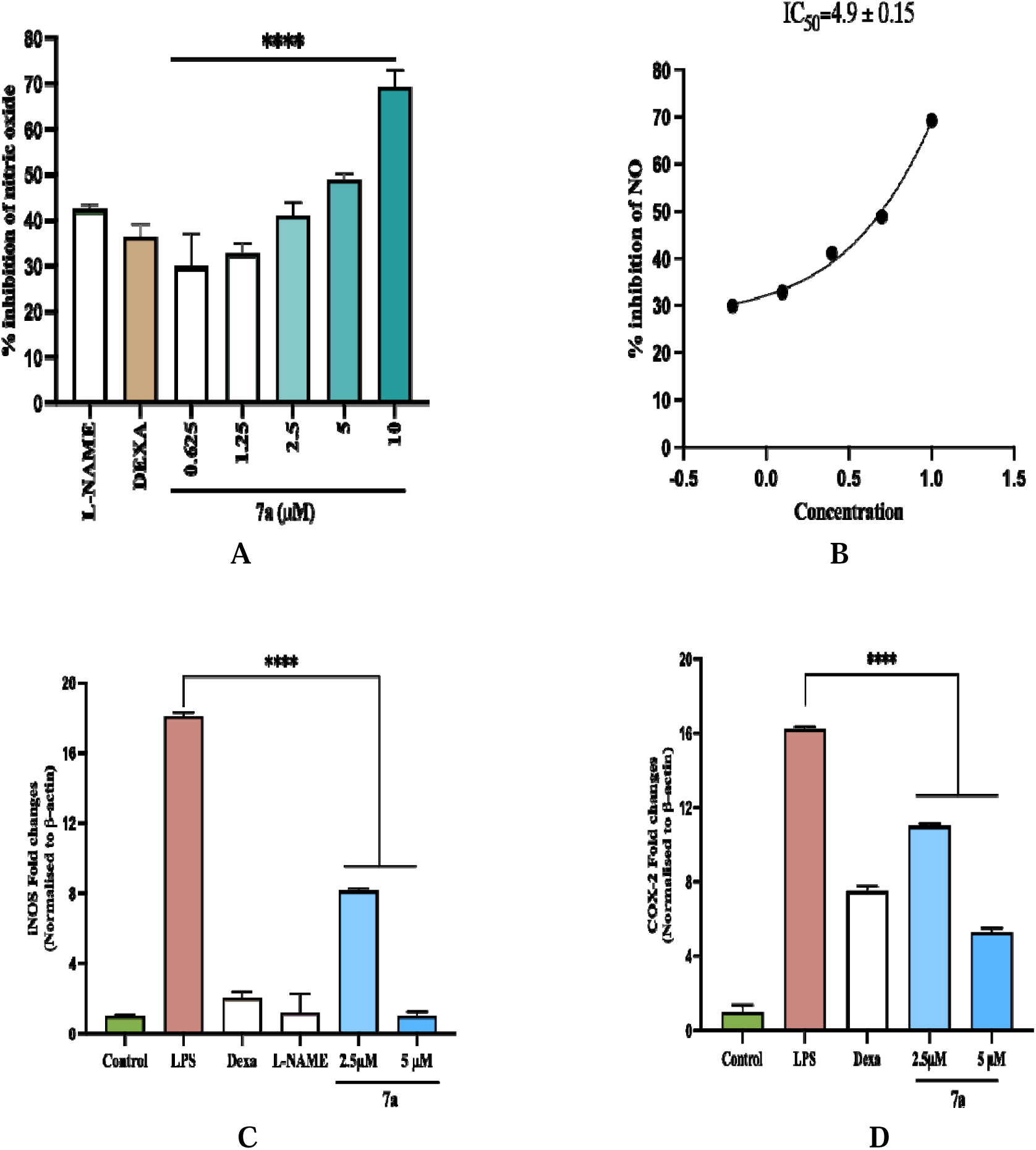
Effect of derivative 7a on LPS-induced RAW 264.7 cells depicting (A) Percentage NO inhibition of 7a. The data were represented as percentage of CONTROL (100%) ± SD (n=3). (B) The IC_50_ graph of the transformed data. The gene expression of (C) iNOS, and (D) COX-2 in RAW 264.7 macrophages stimulated with LPS (1µg/mL) using RT-PCR is represented through the respective graphs. Dexamethasone and L-name were used as positive control. Statistical significance was assessed by one-way ANOVA followed by Dunnett’s test. ** P *<* 0.01, *** P *<* 0.001, **** P *<* 0.0001, vs LPS (1μg/ml).

**Figure 3:**
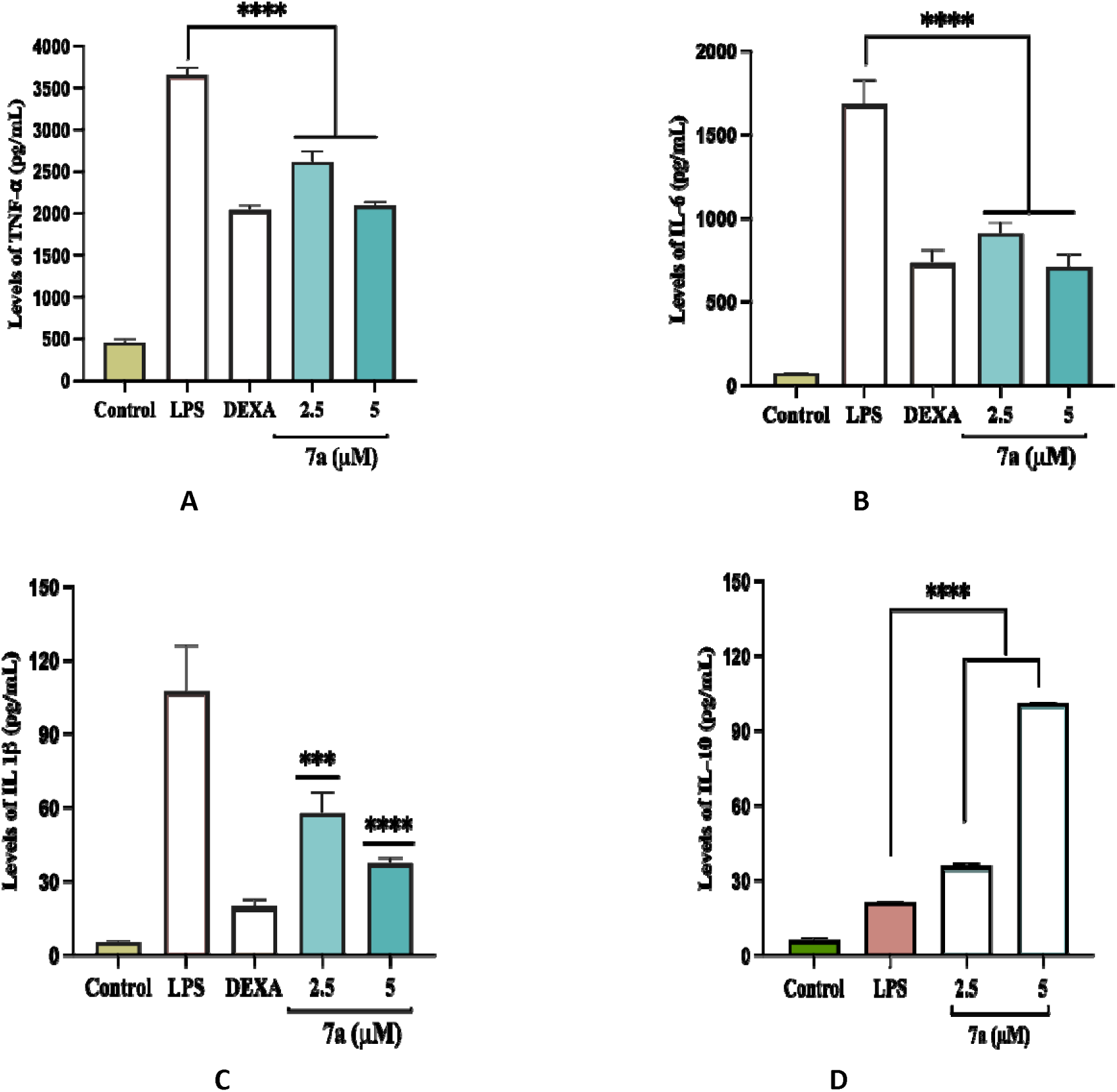
Effect of 7a on the levels (pg/ml) of inflammatory cytokines; (A) TNF-!Z (B) IL-6, (C) IL-1β, (D) IL-10 in pre-treated RAW 264.7 cells stimulated with LPS (1µg/mL) for 24 h. The data were represented as the mean ± SD with n=3. Statistical significance was assessed by one-way ANOVA followed by Dunnett’s test. *** P *<* 0.001, **** P *<* 0.0001, vs LPS (1μg/ml).

**Figure 4:**
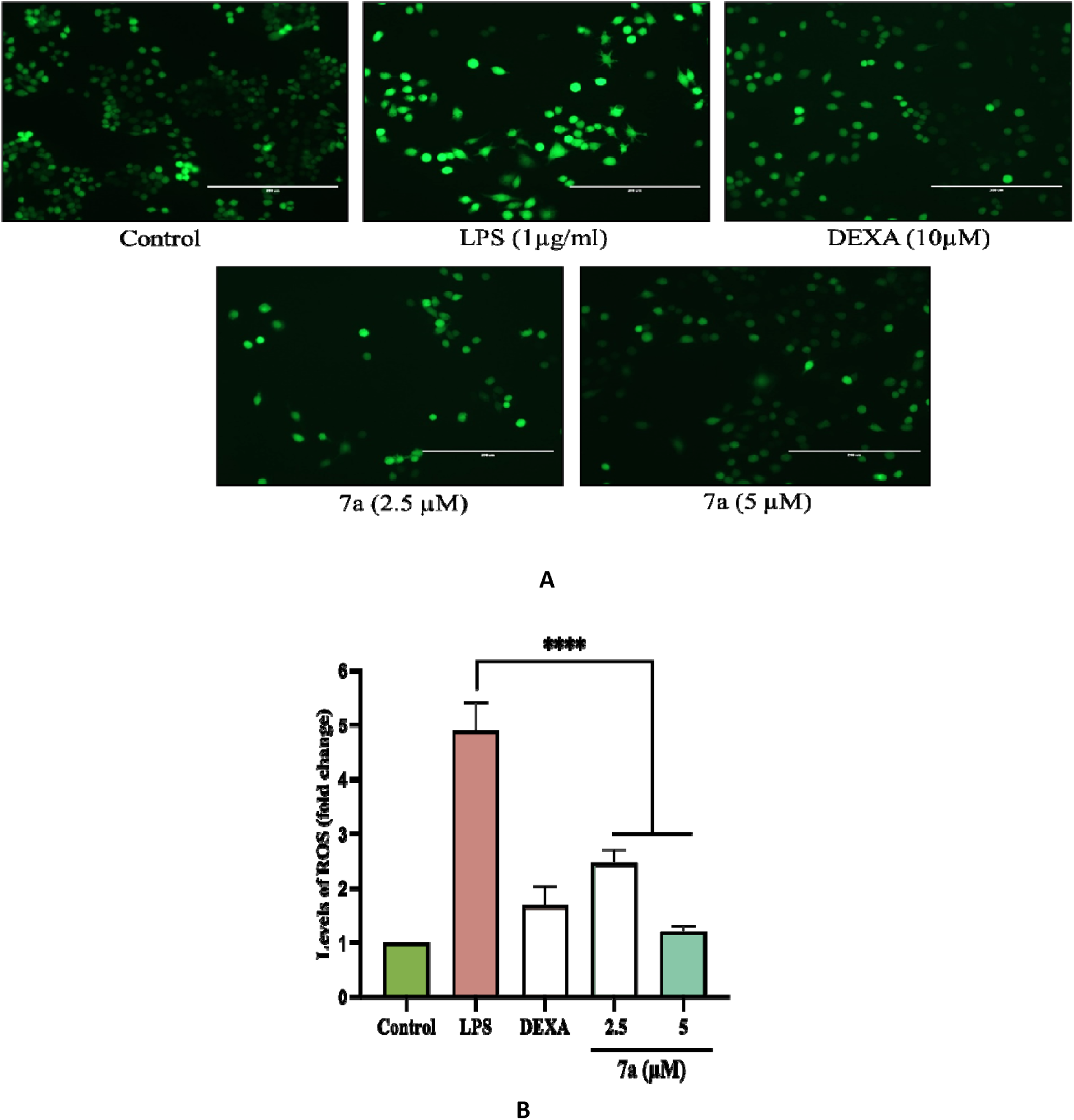
Effect of the derivative 7a on the levels of reactive oxygen species (ROS) in RAW 264.7 macrophages stimulated with LPS (1µg/mL). (A) Assessment of ROS through fluorescence microscopy (20x), (B) Percentage inhibition of ROS analysed by ImageJ. The data were represented as the mean ± SD with n=3. Statistical significance was assessed by one-way ANOVA followed by Dunnett’s test, **** P *<* 0.0001, vs LPS (1μg/ml).

**Figure 5:**
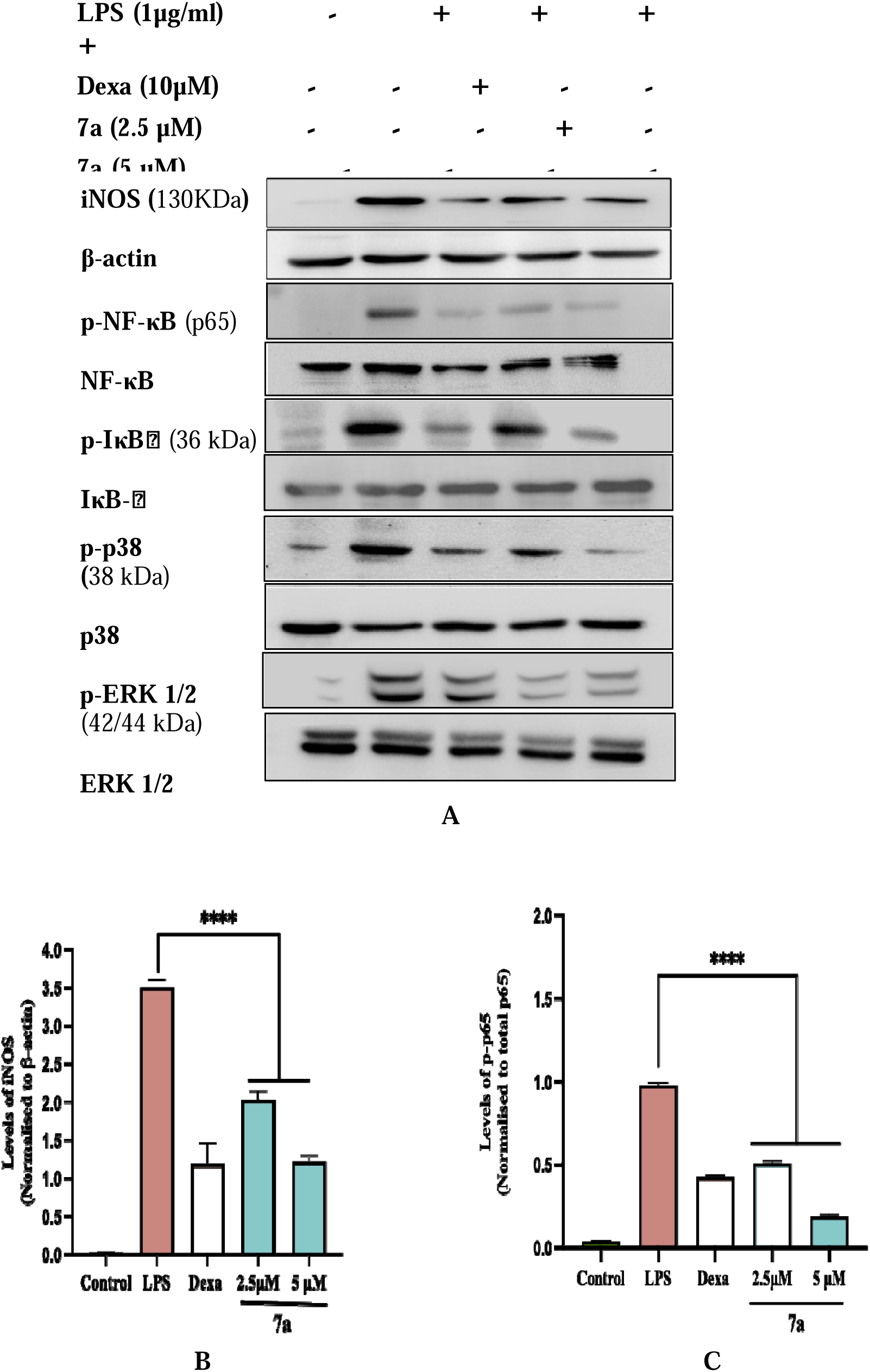
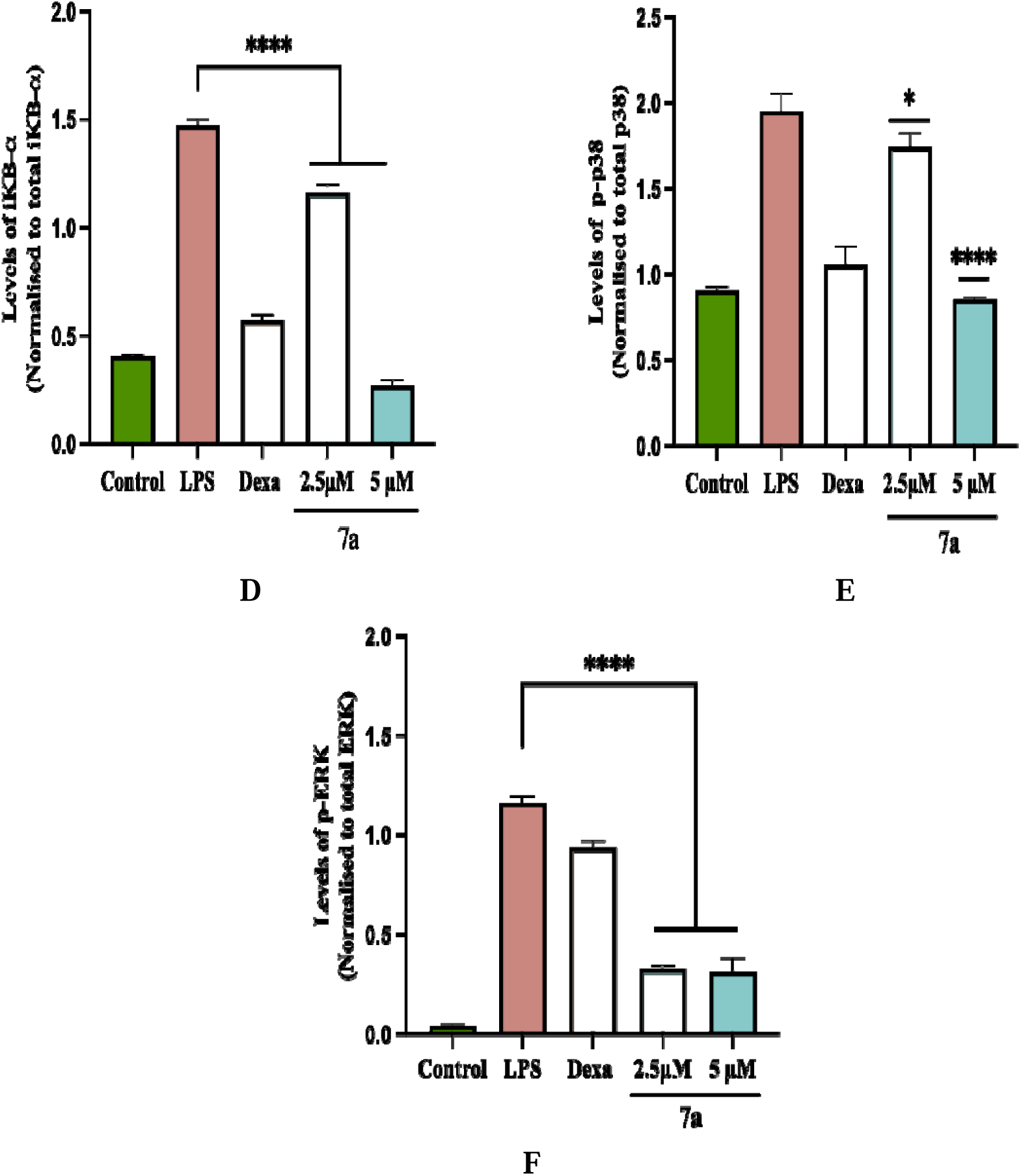
Effect of derivative 7a on protein levels in RAW 264.7 macrophages stimulated with 1 µg/ml of LPS. (A) Western blots of iNOS, p65, IκBl7l, P38 and ERK. Expression levels analysed through ImageJ (B) iNOS, (C) p65, (D) IκBl7l, (E) p38, (F) ERK. The Statistical significance was assessed by one-way ANOVA followed by Dunnett’s test. * P *<* 0.1, **** P *<* 0.0001, vs LPS (1μg/ml).

**Figure 6:**
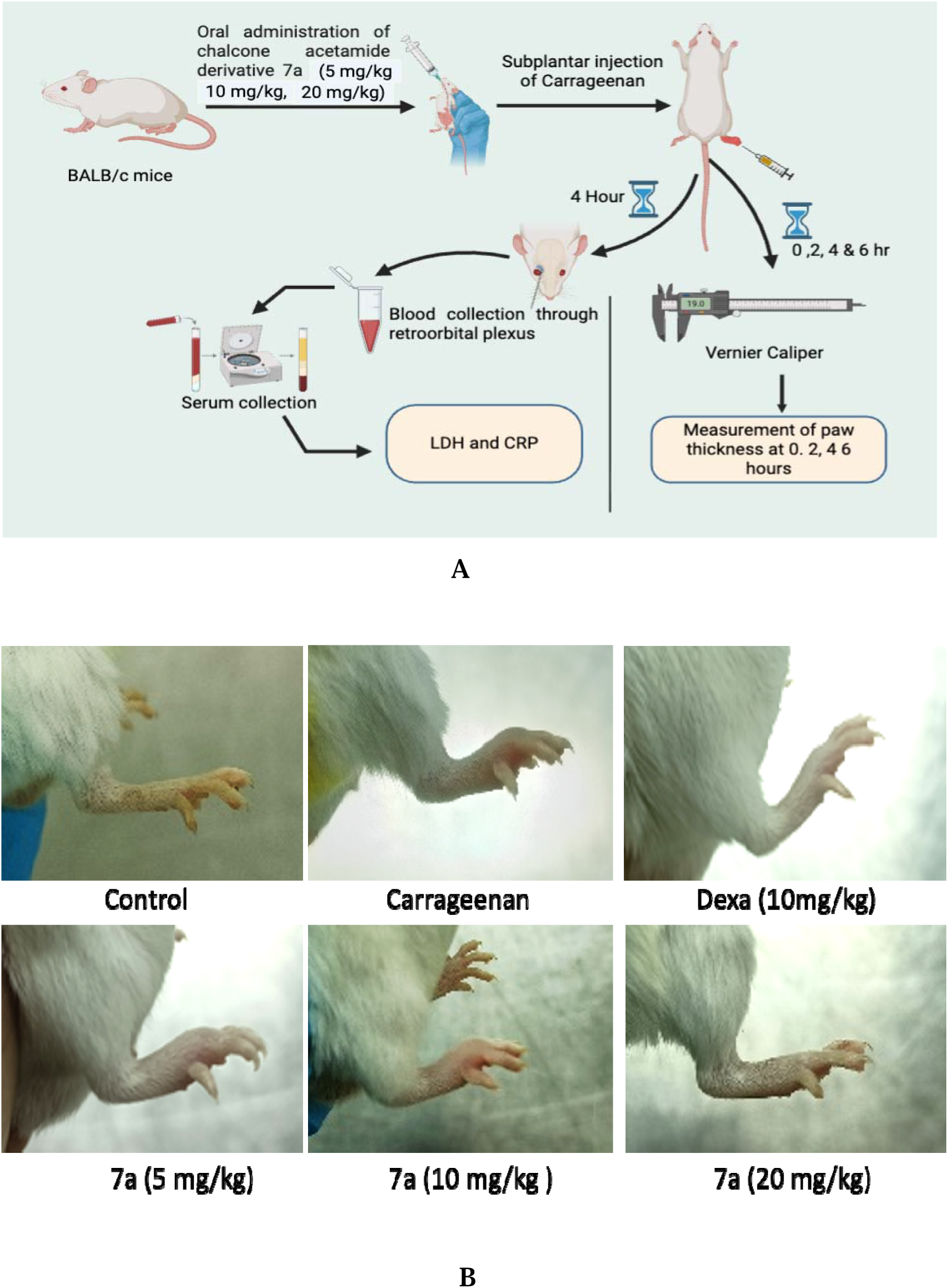
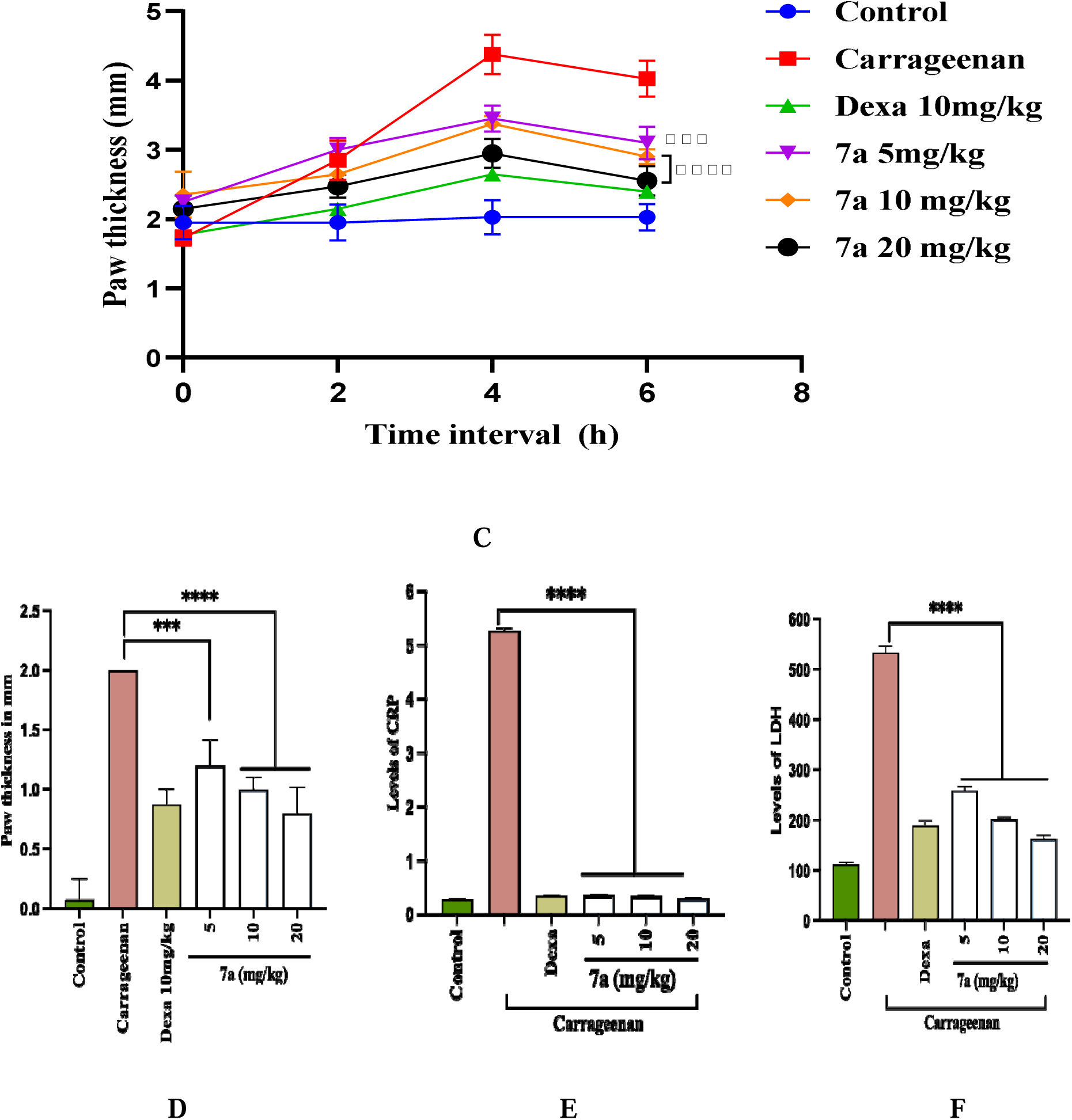
Anti-inflammatory effect of the chalcone derivative 7a in carrageenan induced paw edema model; (A) Schematic illustration of Carrageenan induced paw edema model (B) images of paw depicting control group, carrageenan induction group and 7a treated groups. (C) Graph representing paw thickness of control and treated groups at different time points using vernier calliper. (D) Graph representing paw thickness of different groups at 4 h. The effect of the chalcone derivative 7a on the serum levels of (E) CRP and (F) LDH. The data were represented as the mean ± SD (n=3). Statistical significance was assessed by one-way ANOVA followed by Dunnett’s test. *** P *<* 0.001, **** P *<* 0.0001, vs Carrageenan.

#### 2.3.2 LPS induced systemic inflammation model

Mice were divided into six groups with four animals each; normal control, disease control, Dexamethasone as positive control and test compound 7a treated group (5mg/kg, 10mg/kg and 20 mg/kg). Animals were treated with the compound 7a and after one-hour LPS (1mg/kg) was injected intraperitoneally to induce systemic inflammation. Lipopolysaccharide induces a systemic inflammatory response in mice, leading to elevated levels of proinflammatory cytokines through the activation of immune cells. Nitric oxide and cytokines were measured in the serum after four hours. The nitrite estimation was performed using Griess reagent. The proinflammatory cytokines such as TNF-α, IL-6 and IL-1β were measured using ELISA kits (as shown in figure 7A).

**Figure 7:**
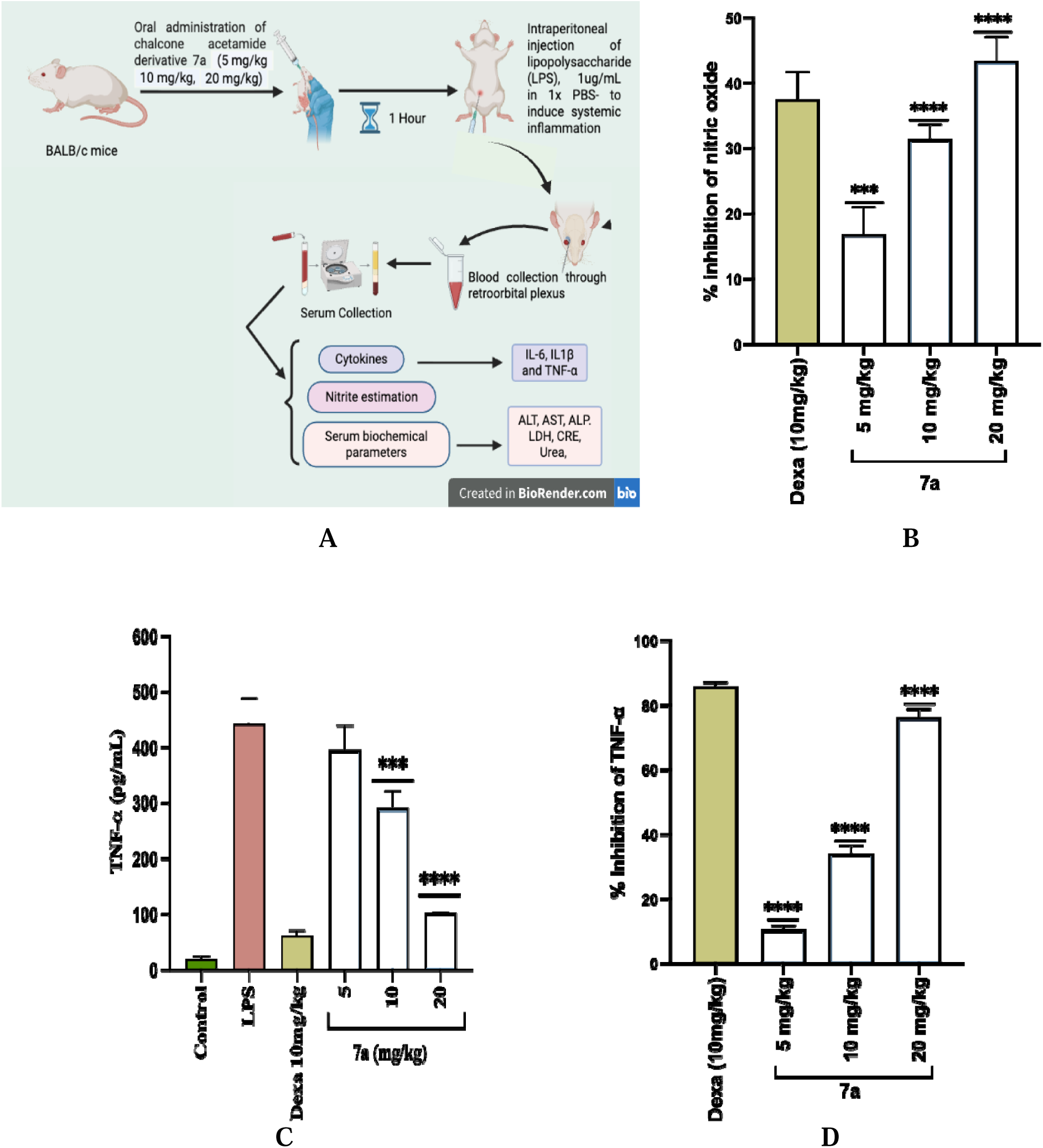
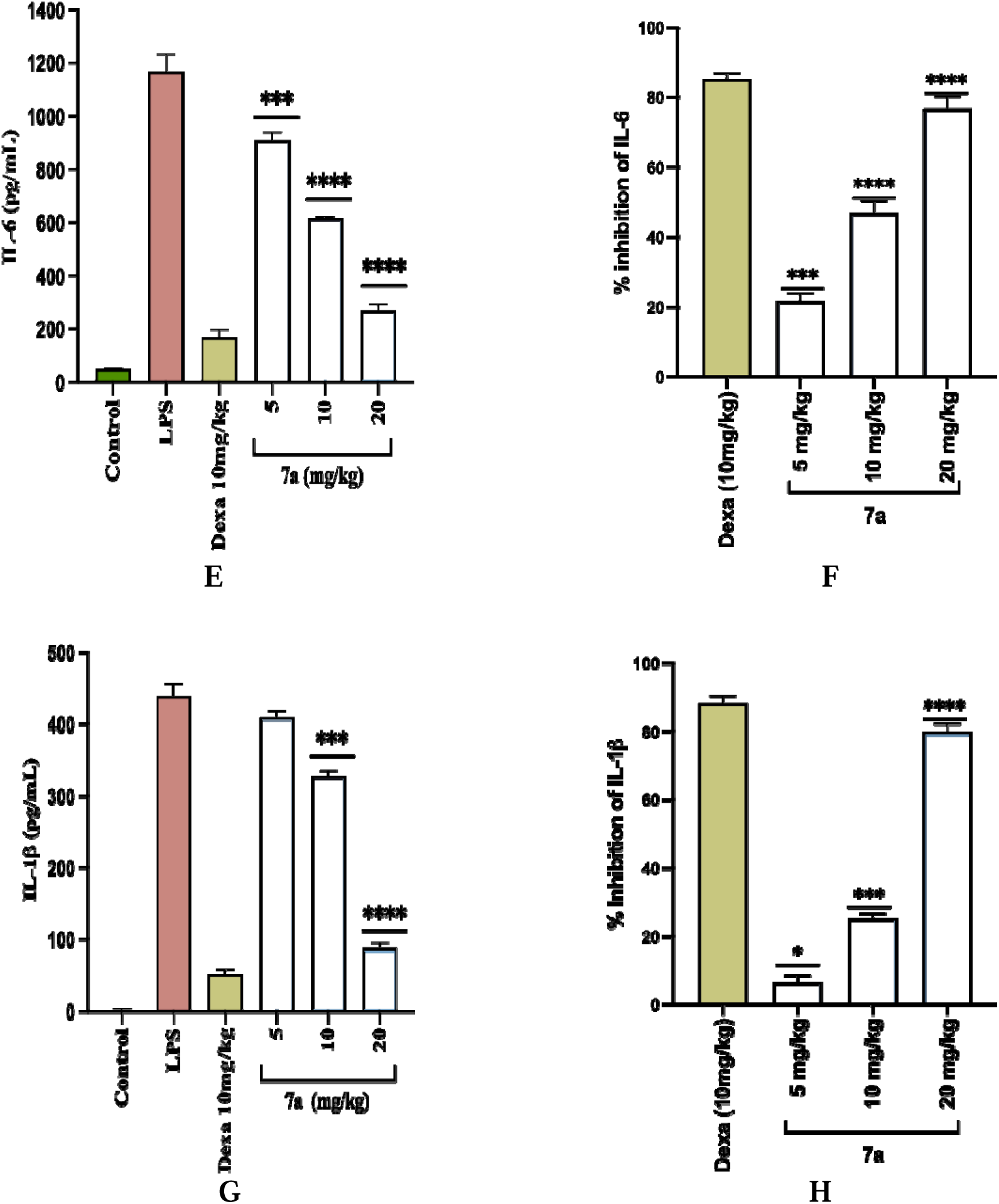
Effect of the chalcone derivative 7a on the pro inflammatory mediators in *in-vivo* LPS induced systemic inflammation model; (A) Graphical illustration of LPS induced systemic inflammation model. (B) Percentage inhibition of NO, (C) TNF-l7l (pg/ml), (D) Percentage inhibition of TNF-l7l, (E) IL-6 (pg/ml), (F) Percentage inhibition of IL-6, (G) IL-1β (pg/ml), (H) Percentage inhibition of IL-1β. Statistical significance was assessed by one-way ANOVA followed by Dunnett’s test. *P *<* 0.1, ** P *<* 0.01, *** P *<* 0.001, **** P *<* 0.0001, vs LPS (1μg/ml).

### 2.4 In-vitro RANKL induced osteoclast differentiation

#### 2.4.1 Preparation of osteoclast precursors from RAW 264.7 macrophages

Murine osteoclasts were generated from RAW 264.7 cells. Briefly, RAW 264.7 cells were plated at a low density in the presence of DMEM with 10% FBS. After 24 h, the media was changed to medium containing compound 7a (2.5, and 5 μM). RANKL (100 ng/ml) was added to induce osteoclastogenesis. Mature multinucleated osteoclasts were seen from day three of RANKL induction. The cells were maintained at 37°C in a humid incubator for 6 days. The supernatant was collected and the inflammatory mediators were measured including NO and cytokines like IL-6 and TNF-alpha.

#### 2.4.2 Tartrate-resistant acid phosphatase (TRAP)-staining

RAW 264.7 cells were used to generate osteoclasts as mentioned above, in the presence or absence of various concentrations of test compound. Briefly, RAW 264.7 cells were plated at a low density in a 6 well plate in complete DMEM. After 24 h, the media was changed to medium containing compound 7a (2.5 and 5 μM) and RANKL (100 ng/ml) was added to induce osteoclastogenesis. After two days, the medium was replaced with fresh medium and cells were maintained in a humid incubator for 6 days. On day 7, the media was removed and cells were stained for TRAP. Cells were washed with 1x PBS and fixed. TRAP staining was done with TRACP and ALP double stain kit (Takara) in accordance with the manufacturer’s recommendations. Osteoclast numbers were quantified by counting the number of TRAP-positive multinucleated cells containing three or more nuclei per field under a light microscope and the images were analysed using imageJ.

#### 2.4.3 ROS estimation

The DCFDA (also known as DCFH-DA), was used to detect intracellular ROS levels. The RAW 264.7 cells were treated with 2.5 and 5μM concentrations of compound 7a and stimulated with RANKL to induce osteoclast differentiation. The cells were maintained for six days while changing media every alternate day. On day seven of the treatment, cells were washed with PBS and incubated with 10μM DCFDA for 20 min at 37°C in the dark. The DCFDA oxidized into fluorescent DCF in the presence of ROS. The cells were again washed with PBS and fluorescence was observed using microscope (Life technologies, EVOS FLOW) and images were analysed through imageJ.

#### 2.4.4 Measurement of inflammatory mediators in RANKL induced RAW 264.7 cells

RAW 264.7 cells were cultured in a 24 well plate (2×10^3^ cells/well). The cells were allowed to acclimatize overnight and further treated with 7a at concentrations of 2.5 and 5 μM and methotrexate (0.0025μM). The cells were then stimulated with RANKL (100μg/ml) and maintained for six days. The supernatant was then collected and evaluated for the detection of cytokines TNF-α, IL-6 and IL-1β, IL-10 by using ELISA kits and nitrite concentration through Griess reaction.

#### 2.4.5 Western blotting

RAW 264.7 cells were seeded at a density of 1×10^6^ cells, in 60mm petri dishes and incubated overnight. Then the cells were treated with different concentrations of derivative 7a and the standard drug followed by RANKL stimulation and maintained for five days for the development of osteoclasts. Then lysates were prepared in RIPA buffer containing a protease inhibitor cocktail, phenylmethylsulphonyl fluoride (2 mM), sodium fluoride (50mM), and sodium orthovanadate (0.5 mM) and analysed for the expression of p65, pIKB-α, P38 and ERK. Protein levels were quantified using Bradford assay and loaded with an equal volume of loading dye. The proteins were then separated using SDS polyacrylamide gel electrophoresis and were electro-blotted onto PVDF membranes, which were blocked for 2 h with 5% bovine serum albumin. The membrane was incubated overnight at 4°C with primary antibody. After that the membrane was washed with 1x TBST and incubated with horseradish peroxidase conjugated secondary antibody. The imaging of the membrane was performed using the ChemiDoc Imaging System (Syngene; Model: GBOX, XX6). Densitometry analysis of the images was conducted using Image J software.

### 2.5. Statistical analysis

The *in vitro* data were expressed as the mean ± standard deviation (SD) of at least three independent experiments. The data management and initial calculations were performed using MS Excel. Statistical significance was determined using ANOVA followed by Dunnett’s test using GraphPad Prism 8 software. p-value of <0.05 was considered statistically significant [^ns^*>*0.05, *p *<*0.05, **p *<* 0.01, ***p *<* 0.001 and ****p *<* 0.0001]. For the *in vivo* experimentation, every outcome was given as mean ±S.E.M., with n = 4.

## 3. Results

### 3.1 Studies on LPS induced RAW macrophages

#### 3.1.1 Effect of chalcone derivatives on cell viability in RAW 264.7 cells

Before examining the anti-inflammatory effect of compounds, an MTT assay was performed to determine the effect of 11 chalcone acetamide derivatives (7a-7k) on the cell viability in RAW 264.7 cells. The cells were treated with different concentrations of the compounds for 48h. Camptothecin was used as a positive control. The results revealed that the derivative 7a maintain cell viability above 90%. Therefore, it was further investigated at concentrations up to 80 µM, where it exhibited no cytotoxic effects, with an IC[[ exceeding 80 µM. In contrast, compounds 7a-7k showed less cell viability at 10 µM concentration. while Camptothecin (positive control) induced 94% toxicity at 10 µM. Consequently, 7a was selected for further assessment of its anti-inflammatory potential in LPS-stimulated RAW264.7 cells.

#### 3.1.2 Effect of 7a on nitric oxide production

Nitric oxide is a key inflammatory mediator produced by inducible nitric oxide synthase (iNOS) [17]. Increased production of nitric oxide can lead to tissue damage and activation of pro-inflammatory pathways [18]. Regulation of nitric oxide generation is therefore an important target for reducing inflammation. Therefore, the effect of derivative 7a on nitrite production was investigated in LPS stimulated RAW 264.7 macrophages and treated with derivative 7a at various concentrations (0.625μM to 10μM). Figure 2A illustrate that 7a showed NO inhibition of 69.25 ± 3.76 % and 48.92 ± 1.28 % at the concentration of 10 μM and 5 μM respectively, indicating that acetamide derivative 7a inhibits excessive nitric oxide production, thereby attenuating the inflammatory response. Moreover, we determined the IC_50_ values and the results suggest that 7a has an IC50 value of 4.9 ± 0.15 μM. Thus, 7a at 5 μM and 2.5 μM was further chosen for a thorough assessment of its anti-inflammatory potential.

#### 3.1.3 Effect of 7a on the levels of inflammatory cytokines

Pro-inflammatory cytokines (TNF-α, IL-6 and IL-1β) are the key regulators of inflammation and play a crucial role in the pathogenesis of various inflammatory and autoimmune disorders [19]. The cytokines production was accelerated in response to the LPS stimulation and contribute to the inflammatory response. RAW 264.7 cells were stimulated with LPS after treatment with the compounds 7a in order to assess its effect on cytokine production The results showed that compound 7a have notably decreased the levels of TNF-α, IL-6 and IL-1β by 42.7 ± 2 %, 57.8 ± 3.1 % and 64.6 ± 5.8 % at 5uM respectively. Besides, it has also significantly fostered the levels of anti-inflammatory cytokine IL-10 (Figure 3D). These outcomes suggested that 7a exhibits better anti-inflammatory effects by inhibiting the levels of pro inflammatory cytokines.

#### 3.1.4 Effect of 7a on intracellular ROS generation

Using the H2DCFDA dye, the impact of the compound 7a on the generation of ROS was studied in RAW 264.7 macrophages. Excessive generation of reactive oxygen species by macrophages indicates oxidative stress and is associated with the onset of multiple inflammatory diseases. The results showed that the test compound 7a significantly inhibited the ROS generation at 2.5 μM and 5 μM concentrations. This implies that 7a has ability to modulate oxidative stress by reducing the production of intracellular ROS as shown in figure 4.

#### 3.1.5 Effect of 7a on the gene expression of iNOS and COX-2

Pro-inflammatory enzymes, including iNOS and COX-2, are crucial in modulating the immune response by promoting the release of diverse inflammatory mediators during inflammatory conditions. Therefore, we evaluated the potential of 7a to inhibit the transcriptional upregulation of inflammation-associated genes in LPS-stimulated RAW 264.7 cells using quantitative RT-PCR. The results showed that 7a significantly suppressed the mRNA expression of iNOS and COX-2 at 2.5 μM and 5 μM concentration, as illustrated in figure 2C and 2D respectively.

#### 3.1.6 Effect of 7a on iNOS, NF-**κ**B and MAPK pathway

To evaluate the effect of 7a on the protein levels of iNOS, p-NF-κB p65, p-IκBα, p-ERK, and p-p38. RAW 264.7 cells were stimulated with LPS, and cell lysates were prepared using ice-cold lysis buffer. Protein levels were analysed through western blotting. iNOS is a pathogenic mediator that plays a key role in inflammation by generating excessive nitric oxide (NO) in response to LPS, a strong inflammatory stimulus. Elevated NO production by iNOS promotes oxidative stress and tissue injury, thereby aggravating chronic inflammatory disorders. The results revealed that treatment with 7a considerably decreased the expression levels of iNOS in a concentration-dependent manner as compared to the LPS treatment. Furthermore, NF-κB and MAPK are central regulators of LPS induced inflammation. As shown in Figure 5, LPS stimulation in RAW 264.7 cells increased the phosphorylation of NF-κB and MAPK pathway proteins. However, treatment with 7a suppressed the upregulation of p-p65, p-IκBα, p-ERK, and p-p38, normalized to their respective total forms.

### 3.2 *In vivo* studies

#### 3.2.1 Effect of 7a on Carrageenan induced paw edema model

Carrageenan is a widely used phlogistic agent in the investigation of acute inflammatory processes and the evaluation of anti-inflammatory drugs. The acute inflammation was induced by injecting 50 μL of carrageenan solution (1%) into the hind paw of mice. The findings demonstrated that carrageenan administration induced a marked increase in paw edema, with maximal inflammatory response observed at 4 hours after injection. However, treatment with derivative 7a have significantly reduced the paw edema and redness in a dose-dependent manner as shown in figure 6B-6D. Furthermore, the levels of C-reactive protein (CRP) and lactate dehydrogenase (LDH) was analyzed in the serum collected after 4h of sub-plantar carrageenan injection. The result showed that derivative 7a effectively reduce the levels of CRP and LDH levels in comparison to the induction group (figure 6B-6F). these findings further validate 7a as a more effective therapeutic compound for the treatment of inflammatory disorders.

#### 3.2.2 Effect of 7a on LPS induced inflammation model

In systemic inflammation, nitric oxide facilitates vasodilation, improving blood flow and supporting the recruitment of immune cells to sites of infection. However, excessive NO production disrupts vascular permeability and cause endothelial dysfunction. Based on the above findings, derivative 7a was further evaluated for its effect against LPS induced systemic inflammation *in vivo* in male BALB/C mice. Wherein, the results demonstrated that the serum levels of NO and pro-inflammatory cytokines were significantly downregulated upon treatment with 7a with the % inhibition of 43.5 ± 3.6, 76.4 ± 2.4, 76.8 ± 3.3 and 79.9 ± 2.4 at the concentration of 20 mg/kg for NO, TNF-α, IL-6 and IL-1β respectively (Fig 7).

Furthermore, blood samples were also collected after 24 hours and the serum levels of biochemical parameters like ALT, AST, ALP, LDH, urea and creatinine were analysed. Increased serum biochemical markers indicate organ dysfunction, with elevated ALP levels suggesting biliary obstruction. Similarly, raised renal markers such as creatinine and urea reflect compromised kidney function, underscoring renal involvement in the systemic inflammatory response. The outcomes suggested that treatment of 7a have significantly lowered the levels of liver and kidney parameters which were otherwise increased due to LPS injection as shown in figure 8.

**Figure 8:**
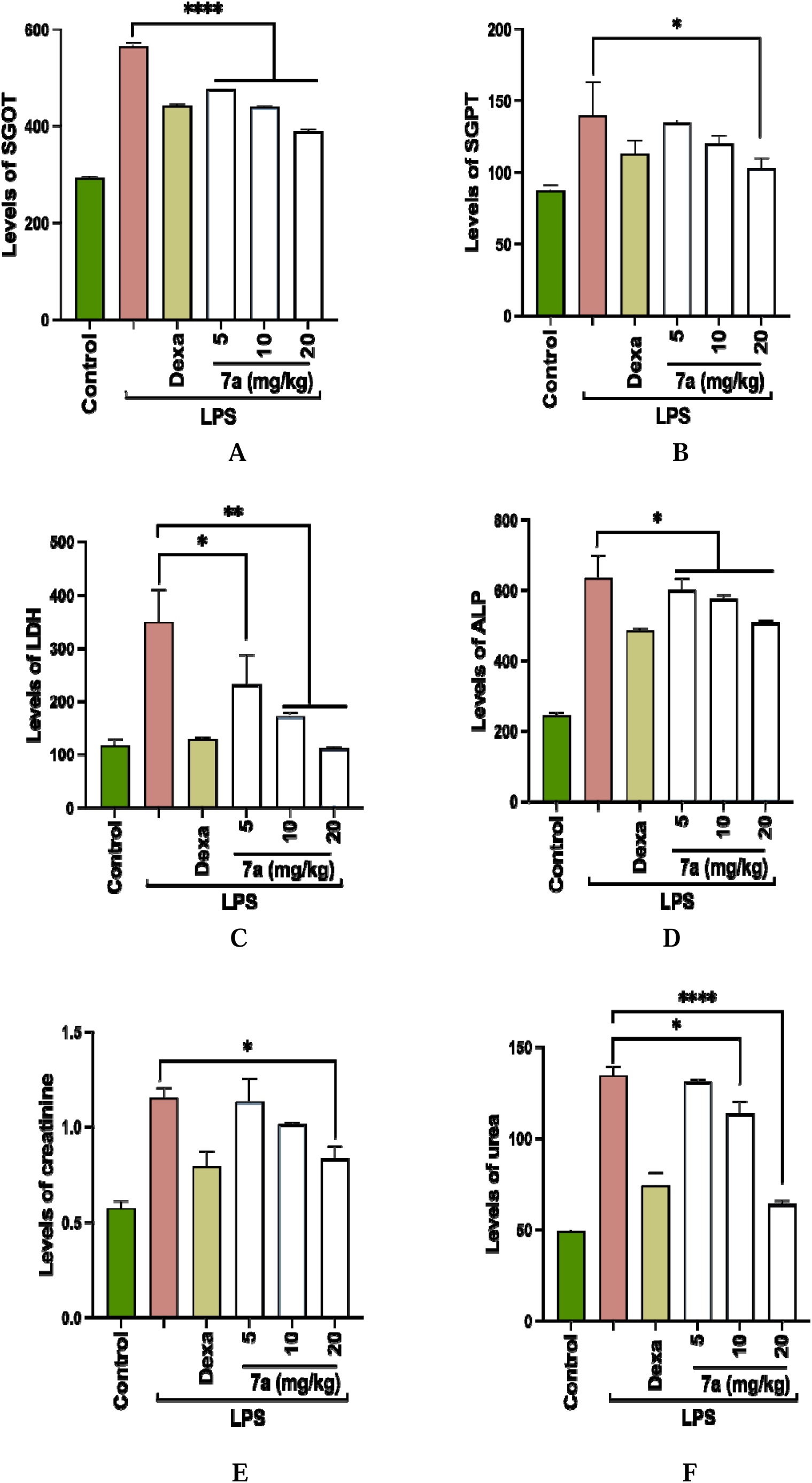
Effect of the chalcone derivative 7a on the levels of biochemical parameters after 24 h of LPS injection; (A) SGOT, (B) SGPT, (C) LDH, (D) ALP, (E) Creatinine, and (F) Urea. Values are expressed as mean ± SD (n=4). Statistical significance was assessed by one-way ANOVA followed by Dunnett’s test. *P *<* 0.1, ** P *<* 0.01, *** P *<* 0.001, **** P *<* 0.0001, vs LPS (1μg/ml).

### 3.3 Anti-osteoclastogenic potential of 7a on RANKL induced RAW 264.7 cells

#### 3.3.1 Effect of 7a on TRAP staining

Tartrate-resistant acid phosphatase (TRAP), serving both as vital histochemical marker and active enzyme, is essential for RANKL-driven osteoclast differentiation and function. Mature, multinucleated osteoclasts exhibit strong TRAP expression, and its activity surges notably during osteoclastogenesis, aligning closely with the emergence of fully developed osteoclasts. In this study, TRAP-positive multinucleated cells were generated by culturing RAW 264.7 cells in the presence or absence of RANKL at the concentration of 100 ng/ml. MTX (0.025 μM) was used as the positive control. The results suggest that 7a decreased the number of TRAP-positive multinucleated osteoclasts in a concentration-dependent manner as illustrated in the Fig 9B and 9C. TRAP-positive mature multinucleated osteoclasts were counted using ImageJ. The highest reduction in multinucleated osteoclast number and percentage area of TRAP positive cells was found at 5 μM concentration of 7a, suggesting its potential anti-osteoclastogenic activity.

**Figure 9:**
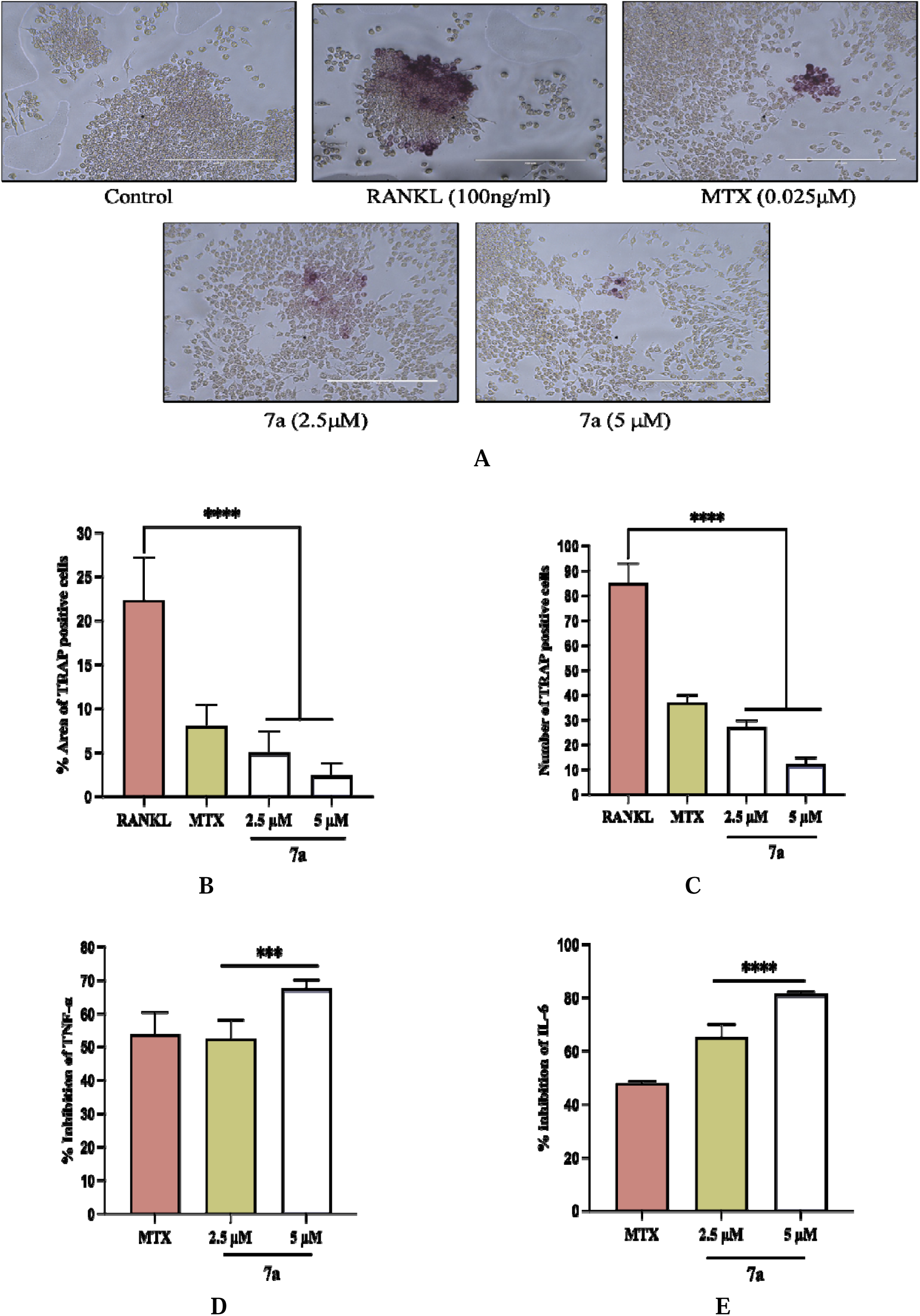
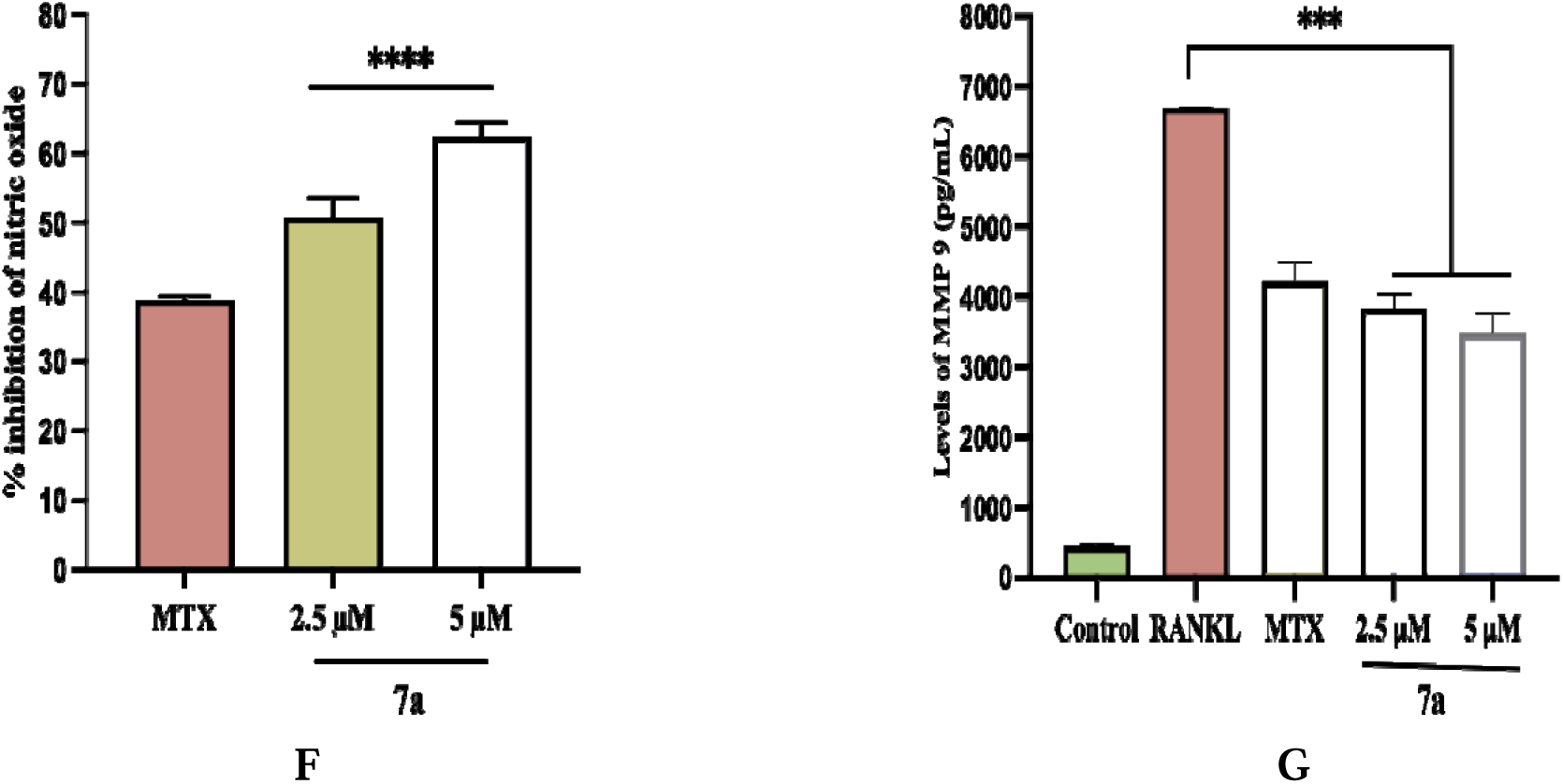
Effect of the selected compound 7a on RANKL induced RAW 264.7 cells; (A) TRAP staining using florescence microscope at 20x. Graph depicting (B) Percentage area of TRAP positive cells, (C) Number of TRAP positive cells. Graphs showing the effect of 7a on different mediators of osteoclastogenesis; (D) TNF-l7l (pg/mL) (E) IL-6 (pg/mL) and (F) Percentage inhibition of NO (G) MMP-9 (pg/mL). Statistical significance was assessed by one-way ANOVA followed by Dunnett’s test, *** P *<* 0.001, **** P *<* 0.0001, vs RANKL (100ng/ml).

#### 3.3.2 Effect of 7a on inflammatory mediators

In this study, we determine the effect of derivative 7a on inflammatory mediators like NO and cytokines including IL-6 and TNF-α. Nitric oxide (NO) exerts a multifaceted influence on RANKL-mediated osteoclastogenesis. RAW 264.7 cells were seeded and treated with 7a at concentrations of 2.5 μM and 5 μM. RANKL (100ng/ml) was added to induce osteoclast differentiation. wherein the results demonstrated that 7a significantly downregulated the levels of TNF-α and IL-6 by 67.5 ± 2.6 % and 81.4 ± 1.0 % respectively, at a concentration of 5 μM as illustrated in figure 9D-9F. These results underscore 7a’s potential to inhibit major pro-osteoclastogenic cytokines, thereby dismantling the cytokine network fueling osteoclastogenesis.

#### 3.3.3 Effect of 7a on the levels of MMP-9

MMP-9 is a member of the matrix metalloproteinase family, a group of enzymes that are involved in the breakdown and remodelling of the extracellular matrix and a complex network of macromolecules that provides structural and functional support to cells and tissues. It plays an important role in osteoclastogenesis, the process of osteoclast formation and activation, which is crucial for bone remodelling and regeneration [20]. The MMP-9 levels were seen in supernatant using ELISA. The results showed that the levels were increased in RANKL treated cells, whereas the group treated with 7a, significantly inhibited the level of MMP-9 (figure 9G). This indicates 7a as a therapeutic candidate for disorders linked to excessive osteoclast activity.

#### 3.3.4 Effect of 7a on osteoclastogenesis via regulating NF-kB and MAPK pathways

This study examined the effect of derivative 7a on the activation of the NF-κB and MAPK signalling pathways induced by RANKL. RANKL stimulation induces IκB phosphorylation, promoting its ubiquitination and subsequent degradation. This process enables NF-κB p65 phosphorylation and nuclear translocation, driving transcription of multiple inflammatory markers, which in turn, were markedly downregulated following 7a treatment. Moreover, RANKL triggers phosphorylation of key MAP kinases—p38, ERK1/2, and JNK—in RAW 264.7 cells. As depicted in the figure 10, The results showed that treatment with 7a effectively attenuated the phosphorylation state of key proteins, such as p-p65, p-IκBα, p-ERK, and p-p38 in a concentration dependent manner, thereby substantially lowering the upregulation of the NF-κB and MAPK pathways.

**Figure 10:**
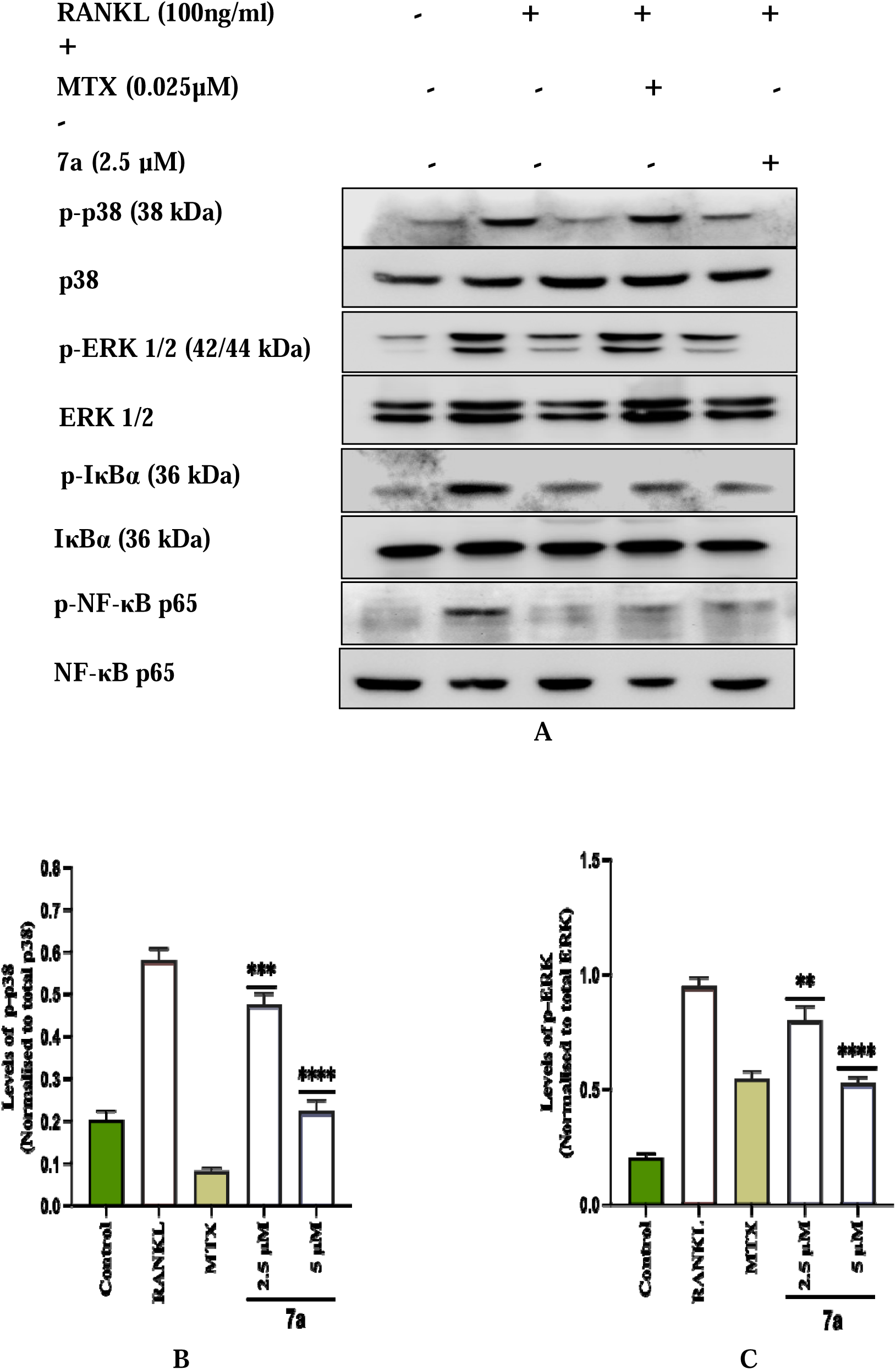
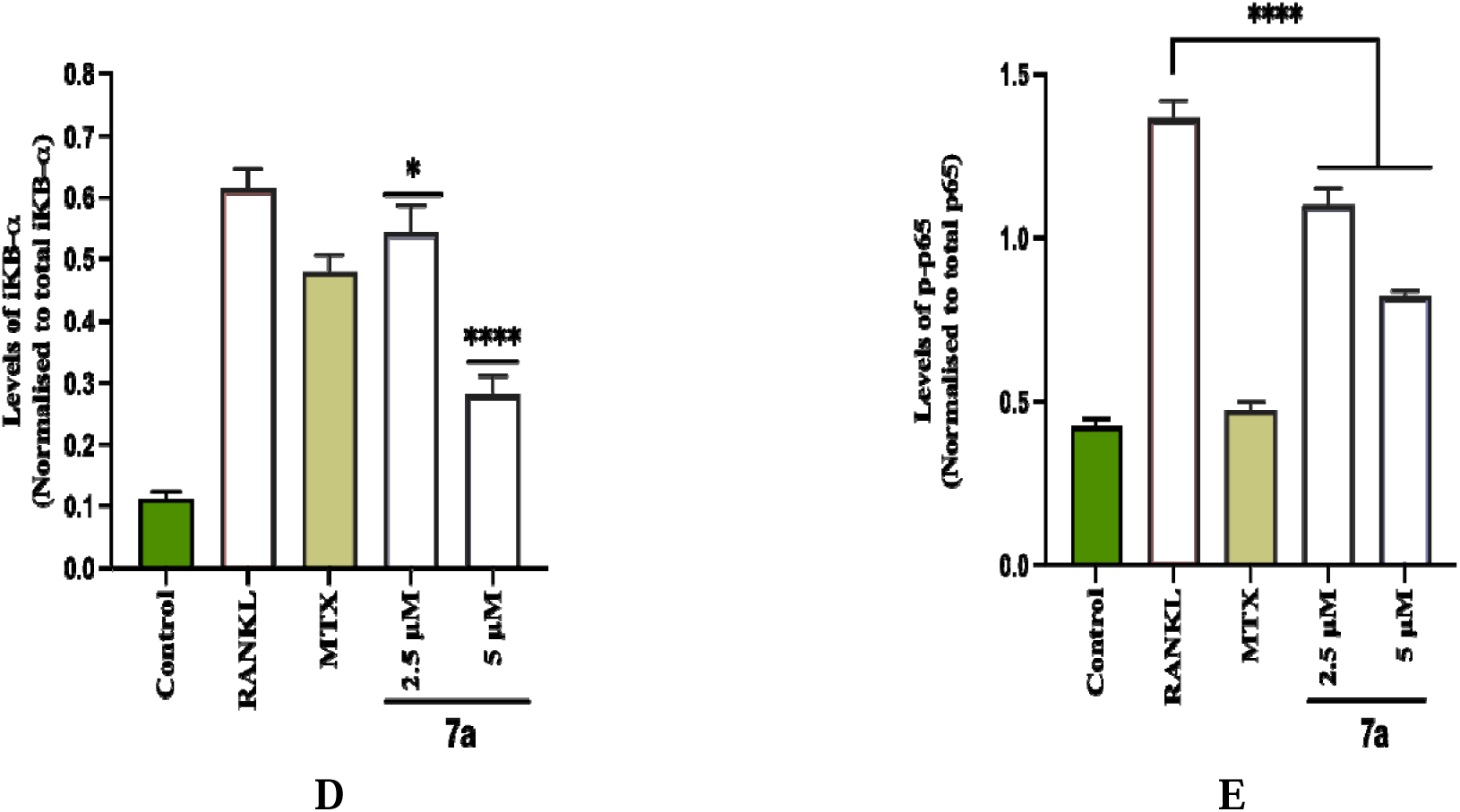
Effect of derivative 7a on p65, IKB-l7l, p38 and ERK expression at protein level, in RANKL induced RAW 264.7 macrophages stimulated with 100 ng/mL of RANKL; (A) Western blots of p38, ERK, IKB-l7l, p65 (RelA). Expression levels analysed through ImageJ (B) p38, (C) ERK, (D) IKB-l7l, (E) p65. Statistical significance was assessed by one-way ANOVA followed by Dunnett’s test, * P *<* 0.1, ** P *<* 0.01, *** P *<* 0.001, **** P *<* 0.0001, vs RANKL (100ng/ml).

**Figure 11:**
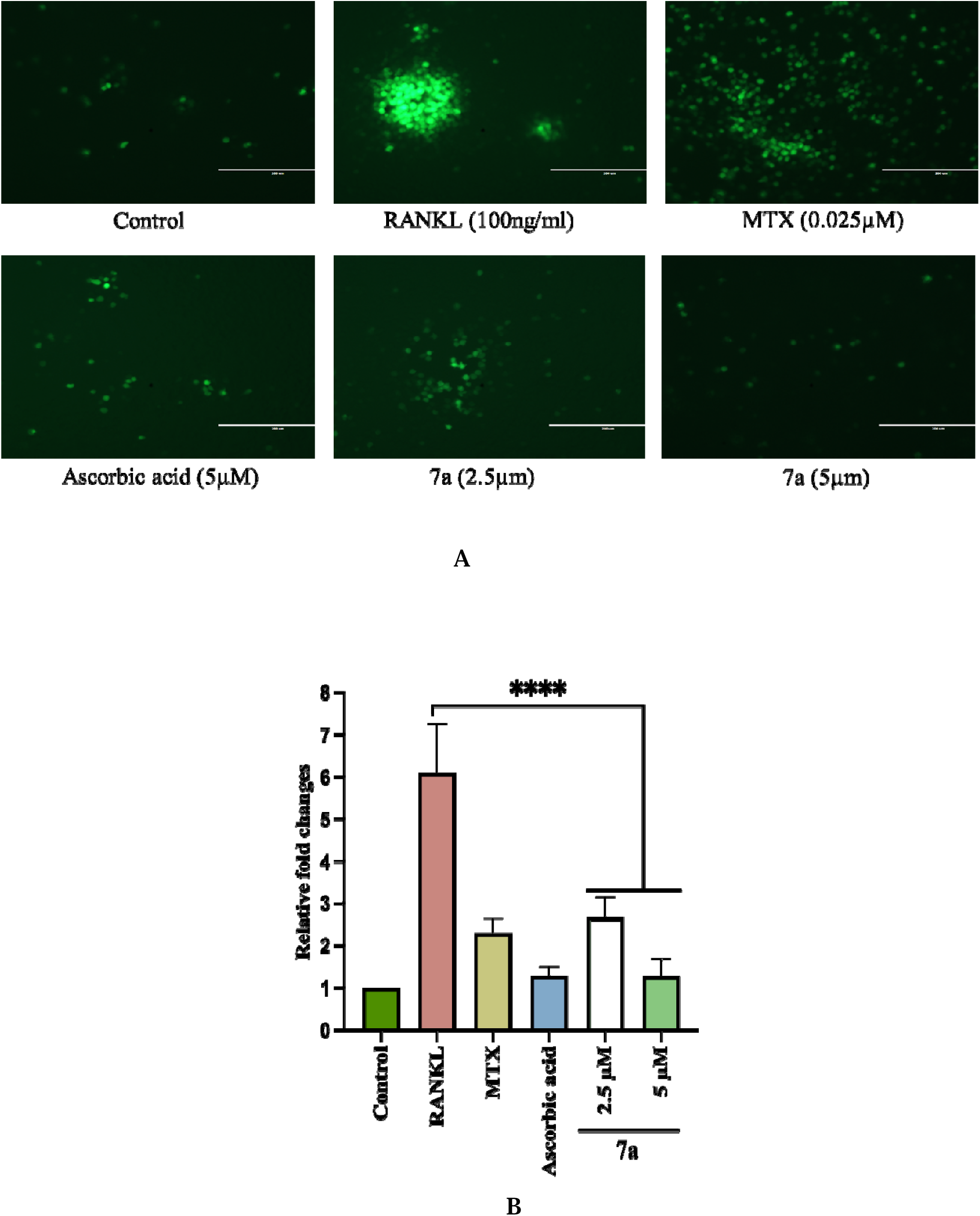
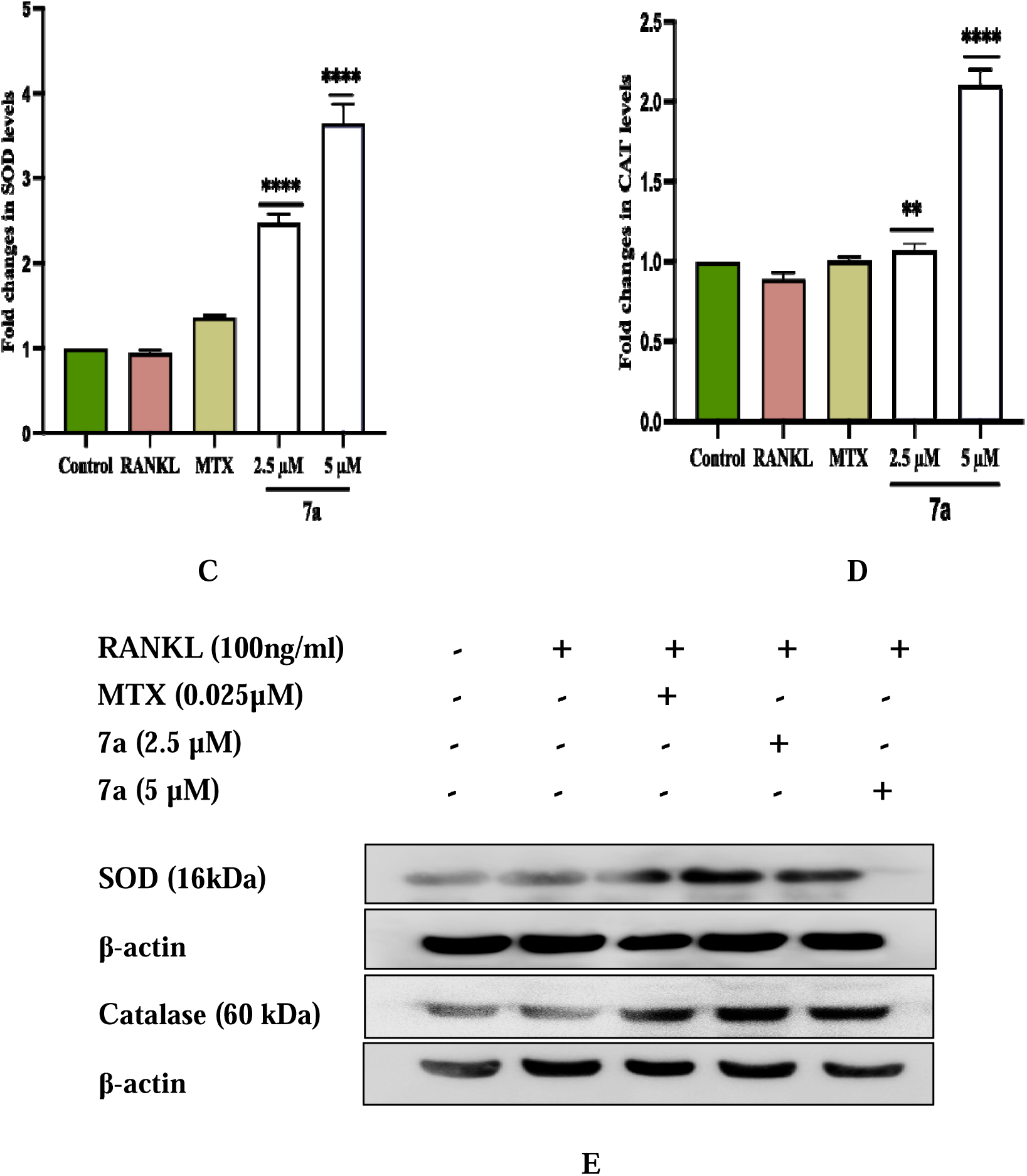
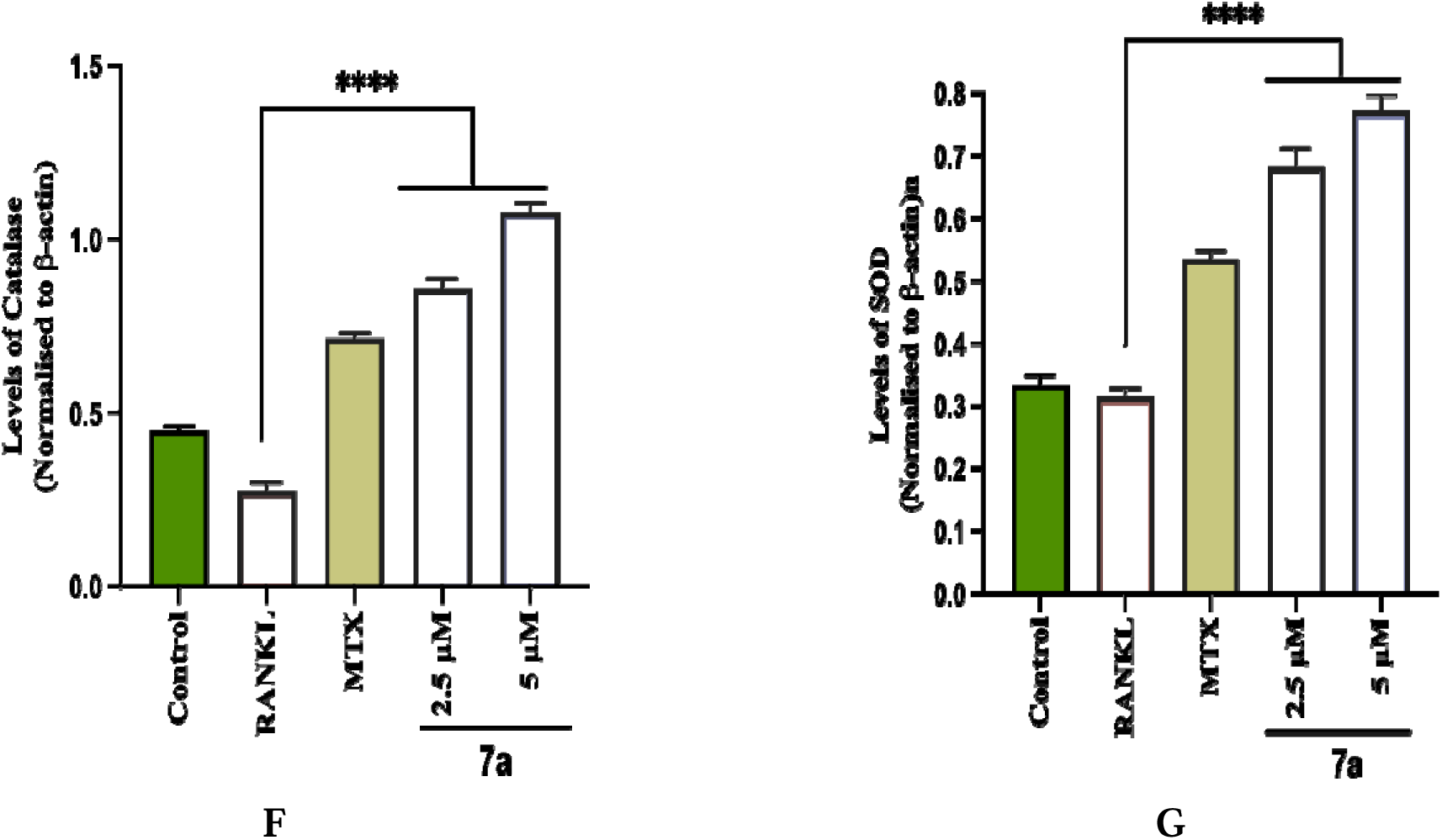
Anti-oxidant effect of the derivative 7a on RANKL induced RAW 264.7 cells. (A) Assessment of ROS through microscopy (20x), (B) Relative florescence of ROS. (C&D) Graph depicting SOD and Catalase levels using ELISA. (E) Western blots of SOD and Catalase normalised using β-actin. (F&G) Protein expression of Catalase and SOD analysed through ImageJ. Data represented as Mean ± SD. Statistical significance was assessed by one-way ANOVA followed by Dunnett’s test, ** P *<* 0.01, **** P *<* 0.0001, vs RANKL (100ng/ml).

#### 3.3.5 Effect of 7a on ROS generation

In this study, we investigated the effect of derivative 7a on RANKL-stimulated ROS production using fluorescent DCFDA dye, wherein the result showed that the intracellular ROS levels were significantly increased on RANKL stimulation, which was subsequently reduced by 7a treatment at a concentration of 2.5 μM and 5 μM respectively. ROS components are essential in the regulation and differentiation of osteoclasts. ROS produced at more than one subcellular site of macrophages was shown to regulate osteoclast differentiation. RANKL is known to induce ROS production in osteoclasts and in turn, ROS can enhance RANKL-induced osteoclastogenesis. Thus, the reduction in ROS levels in osteoclasts induced by 7a may represent a key mechanism underlying the observed suppression of osteoclastogenesis.

#### 3.3.6 Effect of 7a on anti-oxidant enzymes

The regulation of cellular antioxidant enzymes, such as superoxide dismutase and catalase, is a critical aspect of maintaining cellular homeostasis and mitigating the damaging effects of oxidative stress. The effect of 7a on these anti-oxidant enzymes was seen in RANKL induced osteoclasts through ELISA. The results showed that treatment with 7a significantly increased the levels of SOD by 3.8-fold in comparison to RANKL treated group (figure 11C). Additionally, treatment with 7a significantly increased the levels of catalase by 2.3-fold compared to the RANKL treated group (figure 11D). This suggests that 7a can upregulate the expression of these antioxidant enzymes, potentially contributing to its protective effects against oxidative stress-induced bone loss.

#### 3.3.7 Effect of 7a on the protein expression of SOD and catalase

The effect on SOD and Catalase was further validated through western blotting and the results demonstrated a significant increase in the protein levels of these enzymes with 7a treatment. Beta actin was used as a loading control, the concentration of which remained unchanged (figure 11E-12G). The result suggests that 7a efficiently counters RANKL-induced oxidative stress by strengthening endogenous antioxidant pathways.

## 4. Discussion

Inflammation is a highly coordinated biological response that protects the body against harmful stimuli such as pathogens, toxins, and tissue injury. It involves the activation of immune cells and the release of cytokines, chemokines, and signaling molecules that together aim to eliminate the initiating cause and restore tissue homeostasis [21]. While acute inflammation is generally beneficial and self-limiting, persistent or unresolved inflammation can become detrimental, leading to sustained immune activation, tissue damage, and the development of chronic inflammatory and autoimmune disorders such as rheumatoid arthritis (RA) [22]. RA is characterized by continuous synovial inflammation, excessive secretion of pro-inflammatory cytokines, and progressive destruction of cartilage and bone. Increasing evidence highlights the close interplay between immune cells and osteoclasts as a central mechanism driving irreversible joint damage in RA [23]. Non-steroidal anti-inflammatory drugs (NSAIDs) are widely used to alleviate pain and inflammation primarily through inhibition of cyclooxygenase (COX) enzymes and subsequent reduction in prostaglandin synthesis. Despite their effectiveness, prolonged NSAIDs used is frequently associated with serious adverse effects, including gastrointestinal ulceration, renal dysfunction, hepatotoxicity, and cardiovascular complications. These limitations significantly restrict their long-term use and emphasize the need for safer anti-inflammatory alternatives [24]. Disease-modifying antirheumatic drugs (DMARDs), in contrast, are designed to alter the underlying immune pathology of RA rather than simply suppress inflammatory symptoms. Drugs such as methotrexate and other biologic agents target immune cell activation and cytokine signaling pathways involved in disease progression. However, their clinical application is also constrained by substantial side effects, including immunosuppression, increased susceptibility to infections, haematological abnormalities, pulmonary toxicity, and liver damage. Furthermore, DMARD therapy is often costly and does not fully inhibit both inflammatory and osteoclastogenic pathways. These challenges provide a strong rationale for the development of alternative therapeutic strategies that can simultaneously regulate inflammation and bone destruction with improved safety and tolerability [22,25]. These limitations highlight the urgent need for safer therapeutic agents capable of simultaneously targeting inflammatory signaling and osteoclastogenesis.

Chalcones, a class of naturally occurring flavonoids, have emerged as promising candidates because of their broad range of biological activities, particularly their anti-inflammatory properties. Several chalcone-based compounds, including sofalcone, isoliquiritigenin, and xanthohumol, have shown encouraging results in clinical and preclinical studies. However, these compounds suffer from limitations such as poor solubility, variable bioavailability, and metabolic instability [15]. Consequently, continuous efforts are being made to design novel chalcone derivatives with enhanced pharmacological profiles. Previous studies have demonstrated that chalcones can modulate inflammatory signaling pathways and suppress the production of key inflammatory mediators, including cytokines, chemokines, enzymes, and ROS [26, 27]. Importantly, recent studies have demonstrated that chalcone derivatives can also inhibit RANKL-induced osteoclastogenesis and prevent inflammatory bone loss, suggesting their dual therapeutic potential in inflammatory bone disorders [28]. These findings provided the rationale for investigating the anti-inflammatory and anti-osteoclastogenic properties of the chalcone derivative 7a in the present study.

Macrophages play a pivotal role in initiating and amplifying inflammatory responses. Upon stimulation with lipopolysaccharide (LPS), macrophages become activated through Toll-like receptor-4 (TLR4) signaling and produce large amounts of inflammatory mediators such as nitric oxide (NO), prostaglandins, cytokines, and ROS. These mediators contribute directly to tissue damage and perpetuation of chronic inflammation [29,30]. Macrophages play a central role in regulating inflammatory responses and in maintaining tissue homeostasis following injury [31]. RAW 264.7 macrophages therefore represent a well-established *in vitro* model for evaluating anti-inflammatory activity.

In the present investigation, cytotoxicity assessment using the MTT assay confirmed that 7a did not compromise cellular viability, allowing its biological effects to be evaluated without interference from cytotoxicity. LPS stimulation significantly increased NO production and induced the expression of inducible nitric oxide synthase (iNOS), consistent with the established role of NO as a key inflammatory mediator responsible for vasodilation, leukocyte recruitment, and tissue injury [32]. Treatment with 7a markedly reduced NO levels and downregulated iNOS expression at both mRNA and protein levels, indicating effective suppression of macrophage activation. Prostaglandins generated from arachidonic acid via cyclooxygenase (COX) enzymes represent another major inflammatory pathway. COX-2 is the inducible isoform responsible for prostaglandin production during inflammation, whereas COX-3 is considered a splice variant of COX-1 and has been implicated in chronic inflammatory disorders [33,34]. In this study, derivative 7a significantly reduced COX-2 gene expression in LPS-stimulated macrophages, further supporting its anti-inflammatory activity. Cytokines such as TNF-α, IL-1β, and IL-6 are central regulators of inflammatory cascades and play crucial roles in RA pathogenesis by promoting synovial hyperplasia and osteoclast differentiation [22]. Our findings demonstrated that treatment with 7a significantly suppressed the release of these pro-inflammatory cytokines in activated macrophages. Mechanistically, this suppression was associated with inhibition of the NF-κB and MAPK signaling pathways, which serve as master regulators of inflammatory gene expression [35]. These observations are in agreement with previous studies reporting that chalcone derivatives exert anti-inflammatory effects through modulation of NF-κB and MAPK cascades [27]. Together, these *in vitro* findings establish 7a as a potent inhibitor of macrophage-mediated inflammatory responses.

The anti-inflammatory efficacy of 7a was further validated using established in vivo models of inflammation. In the carrageenan-induced paw edema model, oral administration of 7a significantly reduced paw swelling, demonstrating effective suppression of acute inflammatory edema. In LPS-challenged BALB/c mice, treatment with 7a resulted in marked reductions in serum levels of inflammatory biomarkers, including C-reactive protein (CRP) and lactate dehydrogenase (LDH), as well as key pro-inflammatory cytokines such as TNF-α, IL-1β, and IL-6. In addition, 7a normalized serum biochemical parameters including ALT, AST, ALP, creatinine, and urea, which are commonly elevated during systemic inflammation and organ stress. These findings suggest that 7a not only mitigates inflammatory signaling but also provides protection against inflammation-associated tissue injury. Collectively, the in vivo data corroborate the in vitro observations and demonstrate the systemic anti-inflammatory potential of this chalcone derivative.

Chronic inflammation is closely associated with pathological bone resorption through enhanced differentiation and activation of osteoclasts. Osteoclasts originate from macrophage-lineage cells and are primarily regulated by the receptor activator of nuclear factor-κB ligand (RANKL)/RANK signaling axis [36]. Activation of this pathway triggers downstream signaling cascades including NF-κB, MAPK, ROS generation, and transcription factors such as NFATc-1 and c-Fos, ultimately driving osteoclast maturation and bone resorption [37,38]. Tartrate-resistant acid phosphatase (TRAP) is a classical marker of mature osteoclasts, while matrix metalloproteinase-9 (MMP-9) plays a crucial role in extracellular matrix degradation and bone erosion [39,40]. In the present study, RANKL stimulation induced the formation of multinucleated TRAP-positive osteoclasts, confirming successful osteoclastogenesis. Treatment with 7a significantly reduced the number of TRAP-positive cells and suppressed MMP-9 production, indicating inhibition of both osteoclast differentiation and bone-resorptive activity. Reactive oxygen species act as secondary messengers in RANKL signaling and enhance calcium signaling and NFATc-1 activation, thereby promoting osteoclastogenesis [37]. Our results demonstrated that derivative 7a significantly attenuated RANKL-induced ROS generation. Furthermore, 7a enhanced the activity of antioxidant enzymes such as superoxide dismutase (SOD) and catalase, restoring intracellular redox balance. Western blot analysis further confirmed that 7a inhibited phosphorylation and activation of NF-κB and MAPK proteins, highlighting its capacity to suppress critical osteoclastogenic signaling cascades. These findings align with recent evidence linking ROS and MMP-9 to cartilage and bone matrix degradation and underscore the importance of targeting redox-sensitive signaling pathways in inflammatory osteolytic diseases [44,43].

The anti-inflammatory efficacy of the chalcone derivative 7a was systematically evaluated using comprehensive *in vitro* and *in vivo* experimental approaches. Based on its potent anti-inflammatory activity, its effects on osteoclastogenesis were further explored using a RANKL-induced differentiation model. Although the present anti-arthritic assessment primarily focused on osteoclast modulation and bone resorption, the findings provide strong preliminary evidence for the therapeutic potential of this compound. Future investigations should explore its effects in established *in vivo* arthritis models such as collagen-induced arthritis and further elucidate its molecular targets. Overall, derivative 7a emerges as a promising small-molecule candidate with combined anti-inflammatory and anti-osteoclastogenic properties.

## Conclusion

The chalcone derivative 7a demonstrated superior biological efficacy compared to the parent chalcone scaffold. *In vitro*, compound 7a significantly enhanced cell viability and exerted potent anti-inflammatory effects by suppressing the production of pro-inflammatory mediators and reactive oxygen species (ROS) in LPS-stimulated RAW 264.7 macrophages. Mechanistically, this effect was mediated through the downregulation of key signaling pathways, namely NF-κB and MAPK. These findings were further supported by *in vivo* studies, where 7a significantly attenuated inflammation in both LPS and carrageenan-induced acute inflammatory models. Additionally, 7a inhibited RANKL-induced activation of these same pathways in RAW cells, thereby markedly impairing osteoclast differentiation, a critical event in osteolytic and arthritic conditions. Moreover, 7a exhibited strong antioxidant activity, evidenced by reduced intracellular ROS levels and enhanced expression of endogenous antioxidant enzymes, including superoxide dismutase (SOD) and catalase (CAT). Collectively, these results underscore the therapeutic potential of 7a as a multifunctional agent with anti-inflammatory, antioxidant, and anti-arthritic properties. Its dual modulation of oxidative stress and inflammatory signaling highlights its promise as a lead candidate for further development in the treatment of inflammatory and degenerative diseases.

## Declaration of authorship contributions

**Sobia Anjum:** Writing – original draft, Methodology, Investigation, Data curation. **Tazeem Akram:** Methodology, Investigation, Data curation. **Unnatti Sharma**: Investigation, Data curation, **Omesh Manhas:** Data Curation. Investigation, **Jasha Momo H. Anal:** Synthesis, Data curation, **Asha Bhagat:** Supervision, **Zabeer Ahmed:** Validation, Supervision, Resources, Formal analysis, Conceptualization, **Gurleen Kour:** Writing – review & editing, Resources, Formal analysis, Conceptualization, Supervision.

## Declaration of competing interest

The authors declare that they have no known competing financial interests or personal relationships that could have appeared to influence the work reported in this paper.

## Acknowledgement

All authors acknowledge the support provided by the Director, CSIR- Indian Institute of Integrative Medicine, Jammu, India. SA, TA, US, OM acknowledges the financial support in terms of fellowship provided by University Grants Commission (UGC) and Council of scientific and Industrial Research (CSIR), Govt. of India. GK acknowledge the financial support through DST- GAP- 3161(Project file number SRG/2023/001857). Institutional IPR No. is CSIR-IIIM/IPR/00954.

## Abbreviations

ALP: Alkaline phosphate
COX2: Cyclooxygenase 2
CRP: C reactive protein
DCFH-DA: 2′, 7′- di chlorodihydrofluroescein diacetate
DAMPs: Damage associated molecular patterns
DMARDs: Disease modifying anti rheumatic drugs
DMEM: Dulbecco’s Modified Eagle’s Medium
DMSO: Dimethyl sulfoxide
ELISA: Enzyme linked immunosorbent assay
FBS: Fetal bovine serum
IκB: Inhibitor of Kappa B
L-NAME: L-NG-Nitro arginine methyl ester
iNOS: Inducible nitric oxide synthase
LDH: Lactate dehydrogenase
LPS: Lipopolysaccharide
MTT: (3–(4,5,dimethylthiazole–2–yl)–2,5 diphenyltetrazolium bromide
NFATc1: Nuclear factor activator of T cells cytoplasmic 1
NF-κB: Nuclear Factor Kappa B
NO: Nitric Oxide
NSAIDs: Non-Steroidal anti-inflammatory drugs
PAMPs: Pathogen associated molecular patterns PBS phosphate buffer saline
RA: Rheumatoid arthritis
RANK: Receptor activator of nuclear factor kappa B
RANKL: Receptor activator of nuclear factor kappa B ligand
ROS: Reactive oxygen species
SGOT: Serum glutamic oxaloacetic transaminase
SGPT: Serum glutamic pyruvic transaminase
TBST: Tris buffered saline with tween
TNF-α: Tumor necrosis factor- alpha
TRAP: Tartarate resistant acid phosphatases.

## Notes

### Competing Interest Statement

The authors have declared no competing interest.

